# Acoustofluidic Active Flow Sculpting Enables Dynamic, Reconfigurable Cross-Sectional Patterning

**DOI:** 10.64898/2026.05.10.724179

**Authors:** Mehmet Akif Şahin, Seyedmohammadali Hosseinian, Anna Černina, Linus Eing, Daniel Stoecklein, Jinsoo Park, Ghulam Destgeer

## Abstract

Soft materials created with flow lithography exhibit distinct functionality depending on the shape and material composition of the precursor fluids, enabling applications ranging from tissue engineering to anti-counterfeiting. However, current manufacturing techniques largely rely on static nozzle geometries or passive hydrodynamic focusing, which commit to a fixed structure during fabrication and limit the ability to actively reconfigure material architecture in continuous flow. Here, we introduce ActiSculpt, an acoustofluidic platform that replaces physical constructions with dynamic, programmable, non-invasive elements to configure the continuous flow. By exploiting the interplay between laminar stability and acoustic streaming, we decouple deterministic fluid deformation from chaotic mixing, enabling the precise sculpting of cross-sectional flow structures. We demonstrate the high-throughput generation of a diverse library of hydrogel particles with continuously tunable moments of inertia. We further demonstrate continuous fiber fabrication in which acoustic parameters are varied in real time, using temporal control to encode structural variation along the fiber’s length. Finally, we translate this platform into a motion-controlled 3D printer, establishing printability of fully-curable and half-curable fiber configurations and confirming that acoustically programmed cross-sections, along with their capacity for multi-fiber structural assembly, are preserved in the printed output. The result is a platform that overcomes the one-device, one-geometry constraint of existing techniques, enabling not only on-demand reconfiguration between fabrication runs but real-time control of material architecture within a single continuous flow, extending from bench-scale characterization to an operating additive manufacturing process. This spatiotemporal control establishes a new design axis for engineering soft materials with programmable mechanical and functional properties.

## Introduction

In the fabrication of functional soft materials [1–4], 3D bioprinted tissues [5–7], or anti-counterfeiting applications [8, 9], the spatial and temporal architecture of the precursor fluid defines the resolution, heterogeneity, and ultimate utility of the final structure. Ideally, co-flowing precursor streams would serve as sculptable media whose internal interfaces could be shaped and repositioned on demand [10–13]. The physical regime of milli/microfluidics provides the ideal environment for such manipulation: laminar flow precludes chaotic hydrodynamic distortion, diffusion-limited mixing preserves compositional distinctness over time, and interfacial stability protects the integrity of the deformed boundaries [14, 15]. Yet these properties remain underexploited: reliance on static architectures precludes active reconfiguration of fluid morphologies during continuous flow, restricting downstream applications to fixed, predetermined designs [16].

Engineering these fluid architectures has largely relied on passive hydrodynamic manipulation via T-junctions or multi-layer channel designs [17–20]. Strategies such as topological relief patterns [21–25], inertial Dean flow sculpting [26–31], and stacked nozzle configurations [32–34] have demonstrated impressive spatial control over cross-sectional distributions, but share a fundamental limitation: dependence on static, complex microchannel geometries. Intrachannel obstacles (e.g., grooves, pillars, localized constrictions) raise hydraulic resistance, requiring elevated driving pressures that increase the risk of device failure [11], while the added geometric complexity imposes tight manufacturing tolerances, requiring a unique, high-precision architecture for each desired flow profile [35]. This “hard-coded” dependency locks operation into a “one-design-one-function” paradigm, precluding the real-time dynamic programmability required for adaptive material fabrication.

To overcome this rigidity, active strategies, such as electroosmosis, magnetofluidics, acoustofluidics, and optofluidics, have enabled dynamic tunability [36–39]. Among these, acoustofluidics has emerged as a particularly powerful, non-invasive candidate [40–42], traditionally harnessed to manipulate discrete objects such as cells and microparticles label-free [43, 44], or to induce chaotic streaming for rapid fluid mixing or gradient creation [45–47]. However, the use of acoustic forces to deterministically sculpt the interface of continuous co-flows remains unexplored: existing methods offer flow redirection or bulk mixing [48] but fall short of fine-grained, spatiotemporal control over the internal architecture of laminar streams.

In this context, what remains missing is a method to actively reconfigure fluid cross-sectional interfaces in space and time. Such spatiotemporal control holds transformative potential across microfluidics: seamless medium exchange or transfer of cells into wash buffers in biology; dynamic tuning of interfacial areas to regulate reaction kinetics or heat transfer in chemical synthesis; and deterministic sculpting of 3D precursors for barcoding and self-assembly in materials engineering. To bridge this gap, we present **ActiSculpt**, a generalizable framework for active, spatiotemporal flow shaping. The system utilizes localized surface acoustic waves (SAWs) to exert dynamically tunable body forces on co-flowing streams, enabling precise cross-sectional shaping of fluid streams within a simple straight channel. By employing active acoustic fields rather than static structures, our platform decouples fluid manipulation from channel-geometry complexity, so that fluid morphology is defined by the electronic input rather than the physical mold. We validated our approach through rigorous characterization, such as sculpting fidelity versus mixing, batch uniformity, and cumulative fluid displacement. To illustrate the platform’s versatility, we generated shape-coded microparticles and continuous, spatially modulated hydrogel fibers, and further demonstrate translation of this sculpting capability into a motion-controlled 3D printer, establishing an operating additive manufacturing process for acoustically programmed fiber structures.

## Results

### Design and operational principle of ActiSculpt

The ActiSculpt platform integrates modular microfluidics with radio-frequency (RF) control to enable on-demand spatiotemporal sculpting of laminar flows (Figure 1A). The initial fluid architecture is defined by configurable multi-inlets that use hydrodynamic focusing to align streams in either vertical or horizontal stacking configurations, enabling modularity to accommodate additional fluid streams (Figure 1B). These fluids are confined within a simple, straight polydimethylsiloxane (PDMS) microchannel (1038 µm *×* 1017 µm) with one wall covered by a 170 µm-thick membrane on a lithium niobate (LiNbO_3_) piezoelectric substrate. The membrane thickness was optimized to minimize acoustic attenuation while ensuring mechanical stability, with thickness-to-wavelength ratios of 5.84 and 7.58 at the lowest and highest working frequencies, respectively (Figure 1C). Downstream, the active manipulation region comprises four interdigital transducers (IDTs) on the LiNbO_3_ substrate, each with a 250 µm active acoustic wave generation aperture. Unique resonant frequencies, *f*_1−4_ = *λ*_*s*_*/c*, where *λ*_*s*_ and *c* are the wavelength and speed of sound, respectively, i.e., 37, 40, 44, and 48 MHz, are obtained by altering the IDT finger distance, *λ*_*s*_*/*4, by 2 µm, allowing for independent activation. The IDTs are laterally offset to span the channel’s full width (Figure 1D). Upon excitation, the transducers launch Rayleigh surface acoustic waves (SAWs) that couple into the fluid as leaky waves at the characteristic Rayleigh angle (*θ*_*R*_) [49], driving an Eckart’s acoustic streaming flow (ASF) that exerts localized body forces and induces secondary flows in the bulk fluid. Cross-sectional fluorescence images confirm the level of spatial control, showing distinct sculpted flow profiles corresponding to the actuation of specific regions (*f*_1_ through *f*_4_) of vertically stacked co-flow (Figure 1E). Beyond static shaping, the rapid response of the acoustic field also unlocks temporal sculpting: dynamically modulating the IDT actuation signals in real time continuously reconfigures the cross-sectional structure of the bulk fluid, enabling longitudinal, streamwise sculpting, demonstrated in confocal scans of the alternating internal architecture of a continuously extruded fiber (Figure 1F). Together, these elements establish a versatile system for reconfiguring complex fluid morphologies on demand.

**Figure 1:**
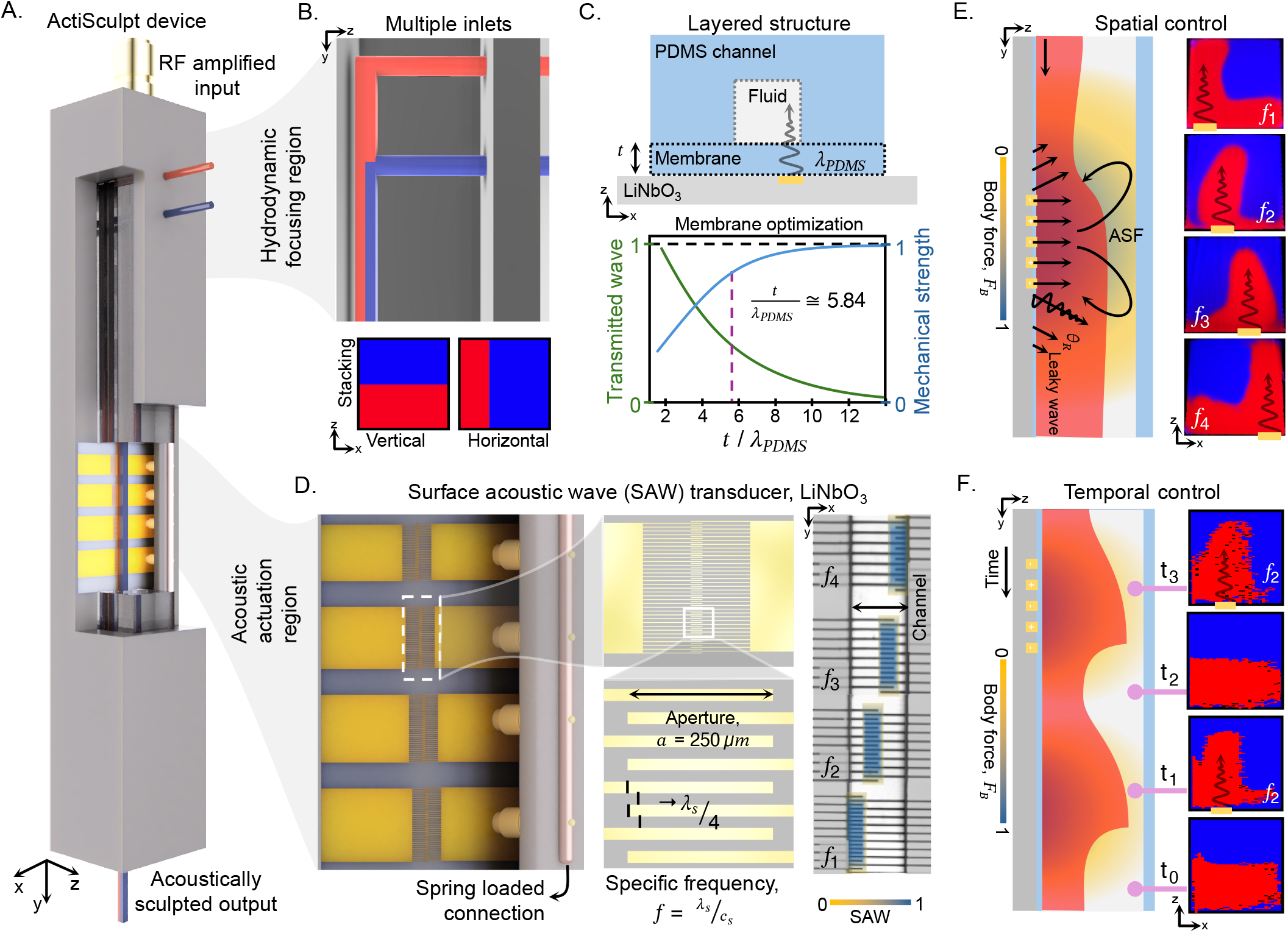
Design and control principle of the modular ActiSculpt device. A. Schematic of the acoustofluidic platform for dynamic flow patterning, integrating two functional regions: a hydrodynamic stacking zone and an acoustic actuation zone for spatial and temporal control of fluid interfaces. B. Stacking orientation is defined *via* inlet design; sequential entry enables vertical stacking, while bifurcation facilitates horizontal layering. C. Schematic of the layered channel structure consisting of a polydimethylsiloxane (PDMS) channel, a membrane, and a lithium niobate piezoelectric substrate. A representative line plot of the transmitted wave and mechanical strength of the membrane along the ratio of the thickness to wavelength provides insight into the optimized selection of the thickness values for the membrane. D. Perspective view of the substrate and the spring-loaded connection with four distinct interdigital transducers (IDTs), each having a 250 µm aperture size and uniquely defined finger spacing and width to correspond to a specific actuation frequency. The inset image shows the overlay of the IDTs, offset from each other to provide full lateral coverage. E. Upon selective excitation of the actuators, Rayleigh surface acoustic waves (SAWs) are generated, which induce leaky waves at the characteristic Rayleigh angle (*θ*_*R*_) within the bulk of the fluid. Resulting localized acoustic streaming flows (ASF), *f*_1_ → *f*_4_, deform the cross-sectional fluid distribution as depicted in fluorescent images of fabricated microparticles, depicting the sculpted flow profile. F. The dynamic modulation of IDT actuation provides temporal control over flow deformation, shown via confocal scan of fabricated fibers. Red color depicts the confocal scan, whereas the blue color is background.

### Integrated platform logic and spectral characterization

To translate acoustic sculpting into fabricated structures via photopolymerization, the microfluidic device was integrated into a custom-engineered experimental setup designed for precise optical and electrical synchronization. A custom holder was fabricated to facilitate alignment between the microchannel, UV mask patterns, and IDTs, ensuring reliable pattern generation and minimizing mechanical drift during polymerization cycles (see Figure S1A). This assembly incorporates spring-loaded pins to maintain robust electrical contact. Central to the operation is the “Active Flow Sequence” control logic, which controls the timing of fluid drive, acoustic excitation, and UV exposure (Figure 2A). Two distinct operational modes were established to exploit this setup, leveraging flow lithographic sequences adapted after extensive literature review [50]: a modified active stop-flow lithography (aSFL) sequence for particle fabrication, where the flow is momentarily halted to “freeze” and polymerize the sculpted cross-section, and a modified active continuous flow lithography (aCFL) mode for fiber fabrication, where acoustic actuation and UV exposure occur simultaneously with sustained flow.

**Figure 2:**
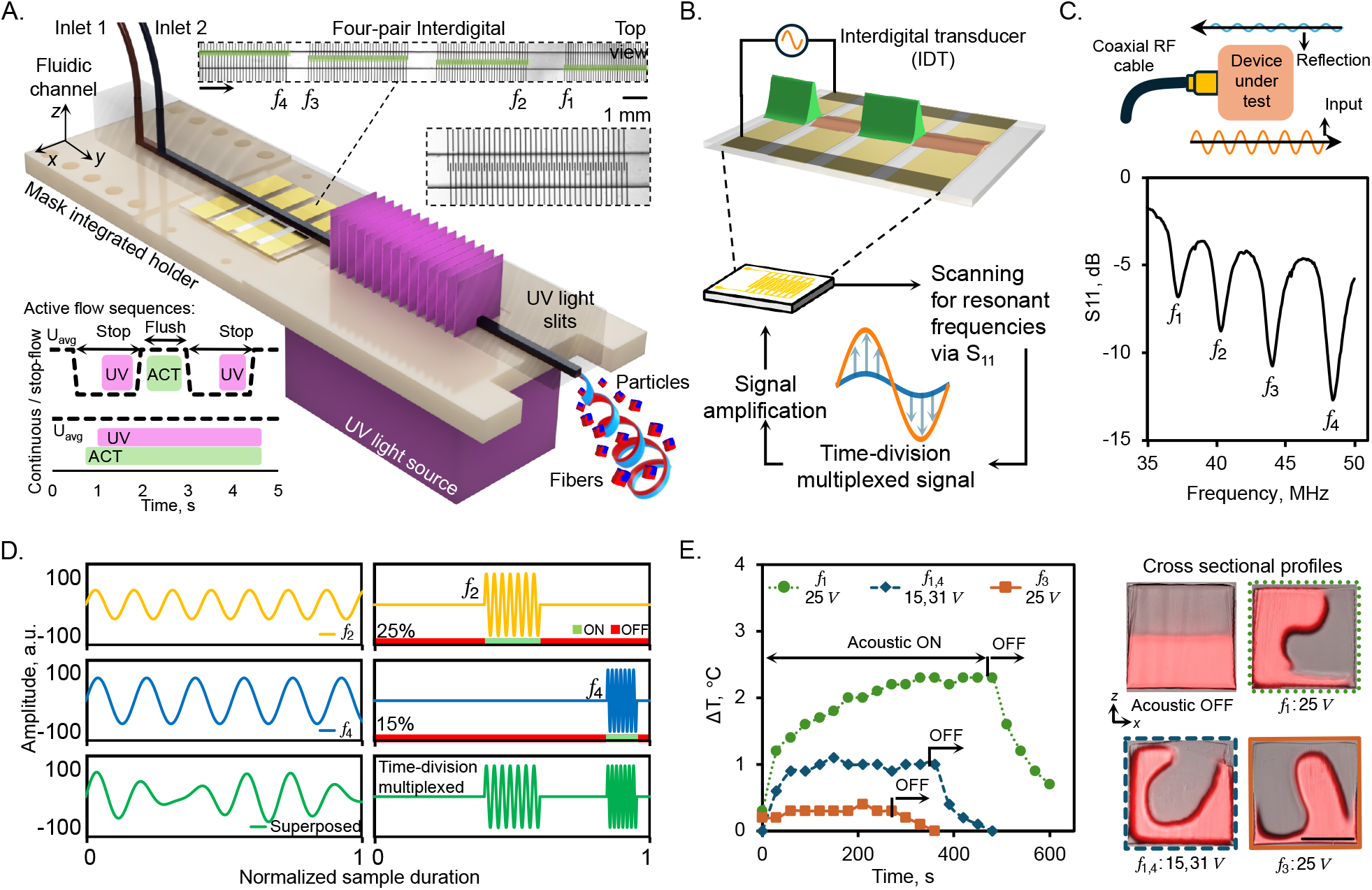
System integration and acoustic control strategies. A. Schematic of the “Active Flow Sequence” logic, detailing the temporal synchronization of fluid pressurization, acoustic excitation, and UV exposure for two operational modes: active stop-flow lithography (aSFL, for particle fabrication) and active continuous flow lithography (aCFL, for fiber fabrication). B. The closed-loop control cycle, illustrating the sequence of spectral characterization (VNA scan), signal definition, and amplification used to maintain optimal acoustic coupling. C. Measured reflection parameter (*S*_11_) spectra, confirming the existence of four distinct, well-separated resonant frequency bands (*f*_1_-*f*_4_). D. Comparison of actuation signal generation methods, demonstrating how the time-division multiplexing (TDM) algorithm preserves waveform fidelity and amplitude integrity compared to the distortion observed with direct signal superposition. E. Real-time differential temperature measurements (Δ*T*) acquired via upstream and downstream thermocouples, confirming that thermal rise remains below 3 °C under optimized TDM duty cycles, ensuring acoustic-dominant flow deformation. Scale bar represents 500 µm.

To achieve precise actuation across the four frequency bands, we implemented a closed-loop control cycle that alternates between spectral characterization and signal amplification modes (Figure 2B). Initially, the transducers are coupled to a vector network analyzer (VNA) to scan the reflection parameter (*S*_11_), which measures the ratio of reflected signal to transmitted signal, providing a strong indication of the natural working frequencies of the IDTs. A relatively lower *S*_11_ value means the same power is used more efficiently to convert electrical signals into mechanical waves. Measuring this parameter is essential: the ~2 µm variation in finger geometry establishes distinct theoretical frequency bands; however, practical factors such as metal electrode fabrication tolerances and temperature fluctuations can shift resonant frequencies. Our device shows *S*_11_ spectra that confirm four well-separated resonances (*f*_1_-*f*_4_: ~37, 40, 44, 48 MHz, respectively), validating the design strategy (Figure 2C).

Once the target frequencies are identified, the system generates a simple sinusoidal function. However, we found that the simple superposition of multiple sinusoidal signals distorted the waveform integrity and weakened acoustic output (Figure 2D). To resolve this, we employed a time-division multiplexing (TDM) algorithm [51] that parses discrete frequencies into non-overlapping time slots. Beyond preserving signal fidelity, this approach introduces tunable duty cycles that provide passive recovery intervals, effectively mitigating heat accumulation. To monitor the thermal environment, we integrated dual K-type thermocouples, positioned upstream (reference) and downstream of the acoustic region, to measure the differential temperature rise (Δ*T*) in real time. Under standard operation, Δ*T* remained below 3 °C (Figure 2E). A comprehensive thermal profile covering more aggressive regimes, including high-amplitude voltage actuation (*>* 130 V), varying flow rate ratios, and extended operation durations, is shown and explained in the Supplementary Information (Figure S2 and Note S1).

### Particle library generation via active stop-flow lithography

To validate the platform’s sculpting capabilities, we employed the active stop-flow lithography (aSFL) sequence to polymerize the fluid’s cross-sectional architecture in mid-flow. The sculpted polymer streams comprise red- and blue-fluorescently labeled polyethylene glycol diacrylate (PEGDA) precursors. Once the precursors are acoustically sculpted, the flow is stopped, and UV light is transmitted through rectangular slits aligned with the channel, forming polymerized particles that “freeze” the deformed cross-sectional flow pattern as red and blue shapes within the microparticle. A random set of collected particles was seeded in a well plate for further image processing, including segmentation and classification (Figure S3 and Note S2). Selective single-actuation of IDTs (*f*_1→4_) generated a diverse library of uniquely “shape-coded” particles, each frequency targeting a distinct lateral portion of the channel, inducing localized acoustic streaming flows that deform the interface between vertically stacked red/blue fluids (Figure 3A). Such deformation is governed by the actuation amplitude: as the feed voltage increases, 10 V to 30 V (plotted left to right in Figure 3), the streaming velocity intensifies, driving a progressive and controlled evolution of the particle morphology from a flat vertically stacked interface to complex, sculpted features (see Figure S4A for high-resolution evolution steps). We further validated the system’s versatility by modulating the initial precursor flow rate ratio from 1:1 to 1:5, applying comparable acoustic actuation to horizontally stacked fluid configurations, and varying precursor viscosities from 4 cP to 57 cP. The input-voltage- and location-dependent flow-deformation trends were retained with expanded sculpting capabilities, resulting in additional unique shape codes (Figure S4B-D).

**Figure 3:**
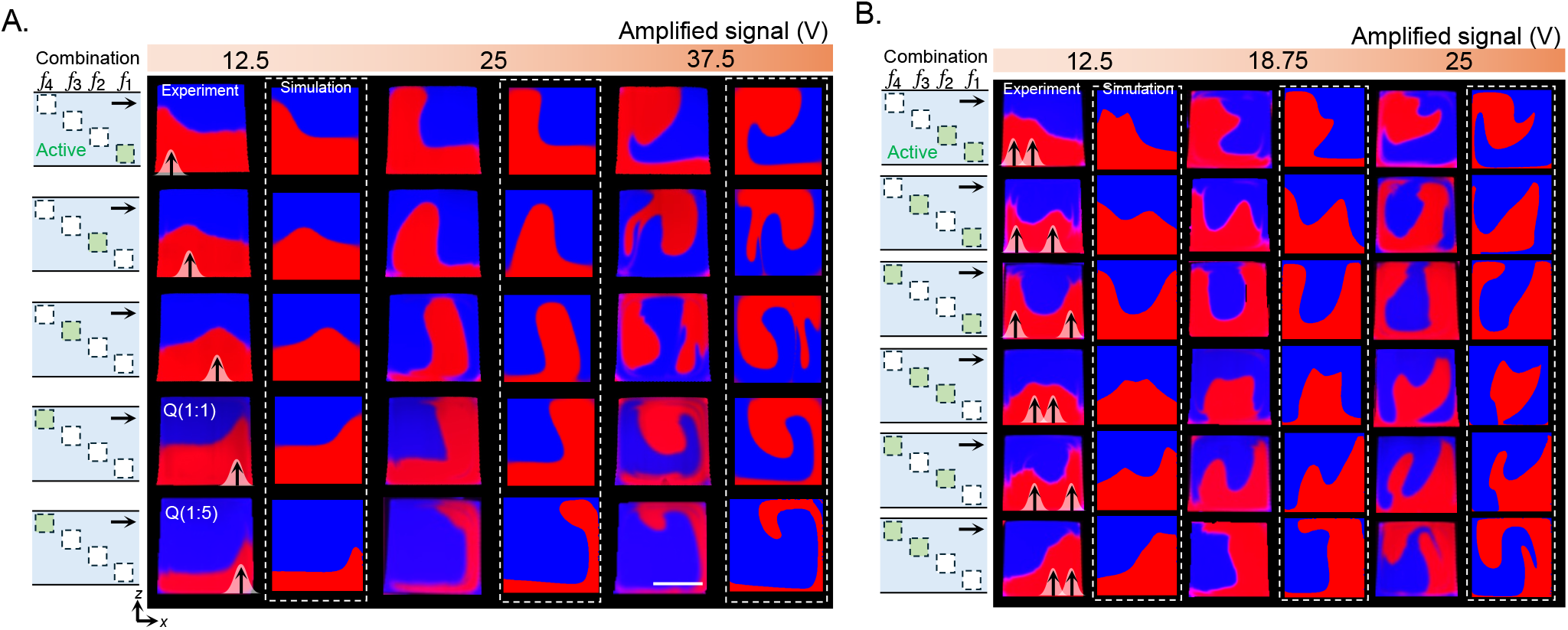
Snapshots of the cross-sectional flow profile captured *via* modified active stop-flow lithography (aSFL). Experimental visualization and numerical simulation of sculpted flow patterns across varying IDT combinations and flow conditions. A. Single actuated IDT output pattern. B. Combinatorial actuations. Amplified voltage (V) increased from left to right (columns) for a given frequency or combination of frequencies (rows), actuating one or two IDTs simultaneously. Scale bars represent 500 µm.

Beyond single-IDT operations, the platform supports simultaneous multi-IDT actuation, enabling multi-point lateral flow perturbations to realize intricate cross-sectional profiles that are not possible with single-IDT actuation (Figure 3B; Figure S4E). Since the transducers are arranged sequentially along the channel, the resulting fluid deformation is cumulative and path-dependent. Consequently, the specific order of activation dictates a distinct sculpted flow morphology; for instance, activating upstream frequencies (*f*_1,2_) yields shapes distinct from those of symmetric counterparts (*f*_3,4_). This flexibility enables advanced operations such as saddle formation (*f*_1,4_ at 15 V), hydrodynamic focusing (*f*_2,3_ at 15 V), and flow rotation (*f*_3,4_ at 15 V). To understand the underlying physical mechanism, we developed a numerical model approximating the acoustic streaming as a distributed body force (Figure S5 and : Note S3). The simulation results (dashed outlines in Figure 3A,B) closely capture the characteristic deformation trends and interface positions, particularly in low-IDT amplitude regimes. At high amplitudes, the model shows a simplified representation of the actual flow structures but still demonstrates good agreement with the experimental data, confirming that acoustic streaming is the dominant mechanism underlying the observed flow sculpting.

### Quantitative characterization of sculpting

While qualitative results illustrate versatility, establishing operational limits requires a rigorous quantitative framework. To achieve this, we developed a rapid image analysis pipeline that automatically segments, filters, and spatially normalizes particle cross-sections from raw fluorescence images to generate statistically significant datasets (see Figure S6 and Note S4 for details). The implemented post-fabrication analysis enabled a robust quantitative assessment of key features, including sculpting fidelity versus mixing, pattern uniformity across batches, cumulative fluid displacement, and structural robustness inferred from the moment of inertia.

### Distinguishing sculpting from mixing

One of the main challenges in acoustofluidics is differentiating controlled deformation of the interface between co-flowing streams from their complete mixing. While acoustic streaming is frequently exploited to accelerate the homogenization of miscible co-flowing streams [52, 53], ActiSculpt seeks to maintain a distinct interface between the fluids (Figure 4Ai). To quantify this trade-off, we analyzed the standard deviations (*σ*) of the red and blue fluorescence intensities within the cured particles, where *σ* ≈ 0.5 represents ideal separation of red and blue segments with zero diffusive mixing and *σ <* 0.05 indicates complete mixing [26, 54] (see Note S5). We established a baseline of *σ* ≈ 0.385 for non-actuated particles (*n >* 80), reflecting inherent diffusive broadening in the laminar flow (Figure 4Ai). We empirically delineated the sculpting regime’ by a threshold of *σ* ≥ 0.25 (half the theoretical ideal), as experimental observations confirmed that the distinct fluid phases remain structurally resolvable above this value. Conversely, values below this threshold (*σ <* 0.25) designate the mixing regime’, where the acoustic streaming forces predominantly drive the homogenization and merging of the co-flowing streams. Upon actuation of *f*_1,4_, the standard deviation exhibited only a moderate reduction to *σ ≈* 0.3, even as amplitudes increased from 0 to 75 V. In contrast, extreme actuation (*V*_*in*_ = 125 V) drove the system toward a mixing-dominant regime, dropping *σ* to 0.18, significantly below the sculpting regime (Figure 4Aii). These results confirm that within the sculpting operational window, the acoustic forces displace the fluid interface without significant chaotic advection, preserving the fidelity of the generated flow shapes.

**Figure 4:**
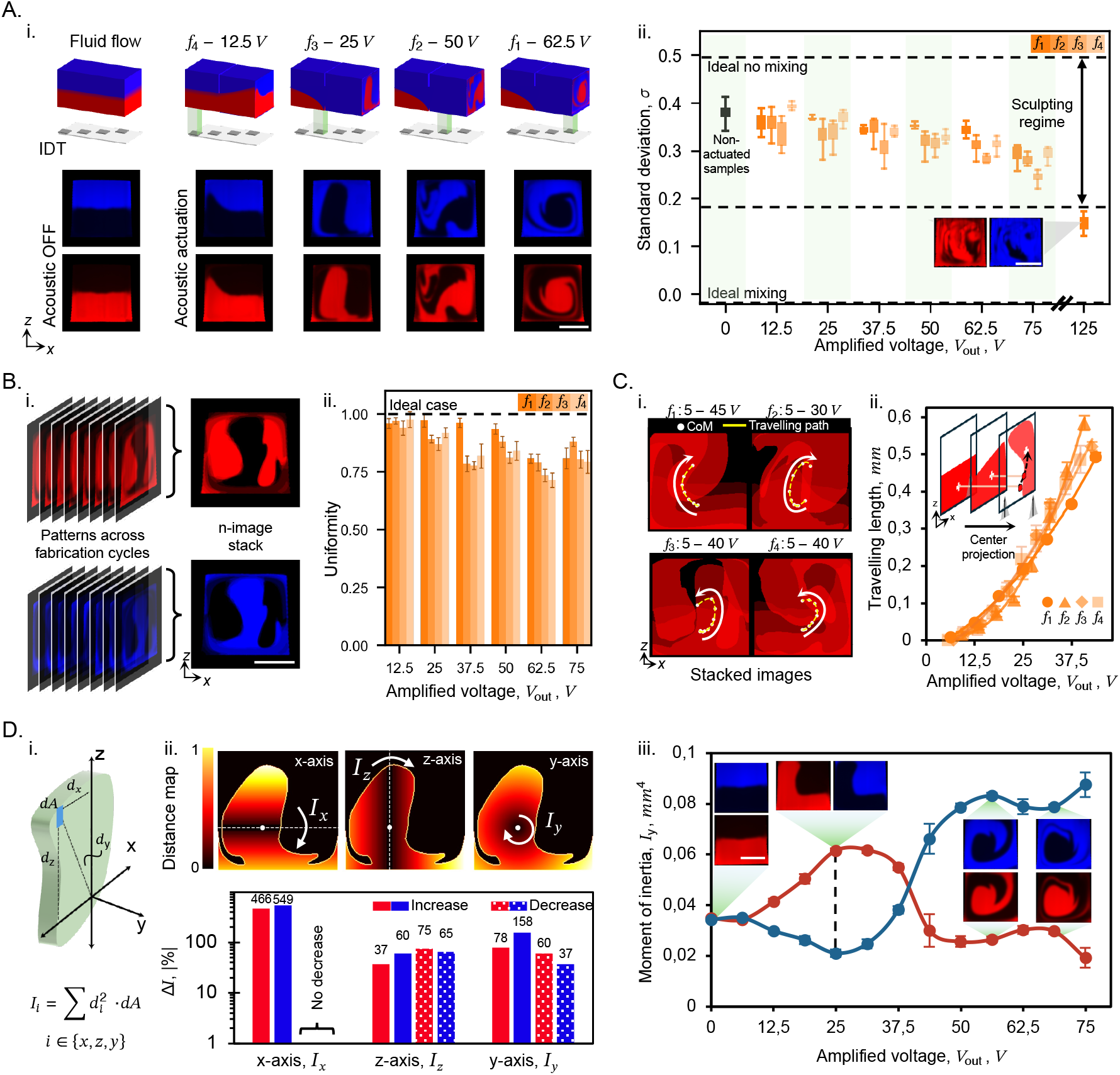
Qualitative characterization of the sculpted flow profiles. A. Dual-color measurements of standard deviation for mixing characterization. i. Examples of non-actuated vs. actuated cases, with varying IDT locations (*f*_1−4_) and input voltages. (top) 3D schematic of the co-flowing red and blue streams. (bottom) Images of cured particles showing the sculpted flow profiles captured in the red and blue fluorescent channels. ii. Standard deviation (*σ*) corresponding to actuation of each IDT (*f*_1−4_) at variable input voltages (*V*_*in*_). Horizontal dashed lines depict the ideal mixing (*σ* ≈ 0.5), experimental reference (*σ* ≈ 0.39), sculpting regime (*σ >* 0.25), and complete mixing (*σ <* 0.05), respectively. (inset) Particle cross-section at high input voltage (*V*_*in*_ = 125 V). B. A uniformity metric, based on pixel-wise overlap of stacked particle images, evaluates pattern consistency across fabrication cycles. i. Stacking the color-separated images to compare pixel occurrence frequency. ii. Bar plot representing the calculated uniformity metric across the IDT actuation at varying *V*_*in*_. Horizontal line shows the ideal case of full similarity. C. Fluid displacement kinematics quantified by tracking the center of mass. i. Stacked images displaying the center of mass for each amplified voltage step; white points are the center of mass, where the dashed yellow lines are a smoothing fit to show the pathway. ii. Traveling path by projection of the center of mass for each corresponding actuation site and voltage level; lines represent a second-order polynomial fit. D. Moment of inertia calculations, derived from pixel-wise mass distribution relative to principal axes, quantify mechanical changes in the sculpted cross-sectional geometry. i. Schematic for the calculation of the moment of inertia on an arbitrary shape. ii. Heat map of the pixel mass distribution (top) and percentage change of the moment of inertia for varying rotation axis for both color channels; patterned bars indicate a decrease. iii. Line plot of *I*_*y*_ for both channels, tracking the variation of the MoI values along the applied voltage, *V*_*out*_. Scale bars: 500 µm (A,B,C), 250 µm (D).

### Process consistency and uniformity

Establishing temporal stability is critical for validating ActiSculpt as a manufacturing method, as thermal accumulation, electrical drift, or mechanical misalignment could degrade flow shaping fidelity over time. To quantify robustness, we developed a uniformity metric based on the pixel-wise consistency of particles pooled from 15 consecutive production cycles of active stop-flow lithography. 10-20 particles were generated per cycle, which were pooled into a single collection vessel for analysis. By digitally aligning a random subset of these particles, we calculated a frequency map of pixel occupancy (Figure 4Bi and Note S6). In this framework, a uniformity score of 1 represents an ideal case where the consensus shape is perfectly preserved across at least half of the population. Empirically, the system demonstrated high robustness; even as acoustic amplitude increased and introduced larger deformations, we observed only a mild decline in uniformity, maintaining a minimum score of ~0.71 (Figure 4Bii). This indicates that even under aggressive sculpting regimes (60 V to 70 V), at least 71%of the pattern area remains spatially consistent, confirming resistance to drift during continuous operation.

### Fluid displacement kinematics

To characterize the kinematics of the sculpting process, we introduced a fluid displacement metric that quantifies the trajectory and magnitude of the cross-sectional transformation. By conducting fine-amplitude voltage sweeps, we tracked the geometric center of mass (CoM) for both the red and blue fluorescent domains at each incremental step. Projecting these spatial coordinates onto the final particle cross-section visualizes the deformation path, in which *f*_1_ and *f*_2_ undergo clockwise rotation, whereas *f*_3_ and *f*_4_ undergo counter-clockwise rotation (Figure 4Ci). The cumulative travel length was then calculated as the summation of Euclidean distances between consecutive CoM positions, effectively capturing the total transport trajectory of the fluid interface. Analysis of these trajectories reveals that the CoM shift is highly responsive to the inner actuation sites (*f*_2_, *f*_3_), while the outer actuation sites (*f*_1_, *f*_4_) exhibit similar kinematic patterns (Figure 4Cii). Nonetheless, all actuation sites achieved a cumulative CoM displacement of at least 0.5 mm within an applied voltage range of 25 V to 43 V. Intuitively, one might expect the outer transducers to induce greater displacement, as they target the near-wall regions where the parabolic flow velocity is lowest. However, because the confined fluid acts as an incompressible continuum, localized acoustic streaming forces propagate globally across the cross-section. Consequently, the observed variations in displacement efficacy between IDTs are driven primarily by frequency-dependent electromechanical coupling and minor acoustic cross-talk, rather than localized flow resistance.

In addition, because the total channel volume is conserved, the kinematic responses of the red and blue domains are inherently coupled. Actuation yields symmetric, anti-correlated shifts in their respective CoM coordinates (ranging *±*100 µm), where a positive displacement in one domain necessitates a corresponding negative shift in the other (Figure S7Ai-iii). The sequenced behavior is also reflected in the decoupled axial displacement plots (x- and z-axes), where the inner actuation sites (*f*_2_ and *f*_3_) reached z-axis saturation and crossed the x-axis zero line at lower actuation thresholds than the outer sites (*f*_1_ and *f*_4_) (Figure S7B). Furthermore, by evaluating the linear deformation regime, we extracted a fundamental kinematic metric: the spatial displacement rate per unit voltage. Across all four actuation sites (*f*_1_ → *f*_4_), the system exhibited an average displacement sensitivity of 15.37 *±* 1.91 µm V^−1^ (Figure S7C; for detailed methodology, see Note S7).

### On-demand control of moment of inertia

The displacement of a structure, *δ*, under loading, *P*, depends on its material stiffness, *E*, and moment of inertia (MoI), *I* [55–58]. The functional implications of cross-sectional flow sculpting extend beyond shape definition to the potential for programmable tuning of mechanical responses, specifically the area moment of inertia (MoI). This parameter, which governs a structure’s resistance to rotational and bending forces, is determined by the spatial distribution of surface area relative to a defined axis (Figure 4Di). Assuming uniform density, we calculated the MoI tensor (*I*_*x*_, *I*_*y*_, *I*_*z*_) for each configuration by integrating pixel-wise distances across the particle cross-section, as depicted in the heat maps in Figure 4Dii (detailed information about the calculation is provided in Note S8). Due to the high aspect ratio of the initial rectangular pattern, the system exhibited pronounced sensitivity in *I*_*x*_; even minor acoustic perturbations disrupted the flat baseline, inducing up to a 5.5-fold increase in this area moment of inertia. In contrast, the remaining moments displayed broad, asymmetric tunable ranges across the experimental dataset, scaling from −37% to +158% for the polar moment (*I*_*y*_) and from −75% to +60% for the transverse moment (*I*_*z*_) (Figure 4Dii, Figures S8 and S9). Another aspect of the modulation is its continuous, yet non-monotonic, nature. For instance, increasing the acoustic amplitude of *f*_1_ initially raises the *I*_*y*_ of the red domain from a baseline of 0.031 mm^4^ to ~ 0.056 mm^4^, before driving an area redistribution that sharply reduces it to 0.013 mm^4^ at higher power levels. A corresponding inverse trend was observed in the adjacent blue domain (Figure 4Diii): the initial value of 0.032 mm^4^ was first reduced to 0.019 mm^4^, then ramped up to 0.082 mm^4^, confirming coupled, dynamic modulation of internal architecture toward targeted geometric behaviors.

### Elongated fiber fabrication

Having established the quantitative framework for cross-sectional sculpting, we extended the ActiSculpt platform from discrete particle generation to continuous fiber manufacturing. By replacing the discrete-pattern photomask with a single-slit geometry and transitioning from the active stop-flow sequence to active continuous-flow lithography (aCFL), the system can sustain the extrusion of hydrogel fibers (Figure S1B). This operational shift leverages the same acoustic architecture but requires precise management of the polymerization kinetics to maintain flow stability. To facilitate smooth extrusion against accumulated back pressure due to polymerized material, we utilized the oxygen-inhibition dead zone’ inherent to the permeable PDMS walls; this unpolymerized lubrication layer acts as a slip boundary, preventing adhesion and allowing a polymerized fiber to glide through the channel. Optimization of the UV exposure parameters revealed a critical trade-off between mechanical integrity and processability: while stiffer fibers resisted deformation, they increased hydraulic resistance and clogging risk. Consequently, the UV exposure window was strategically positioned near the channel outlet to minimize the frictional path length, while the UV exposure was dynamically balanced with the flow rate to ensure the fiber exited the device in a fully cured yet pliable state. The fabricated fibers were then imaged by confocal microscopy to visualize their cross-sectional patterns (see Note S9 for details).

### Fiber fabrication with static actuation

To validate the stability of the continuous extrusion process, we first generated fibers under constant acoustic actuation (*f*_2_ at 40 V) and without actuation as a reference (Figure 5Ai). We used confocal microscopy scans to reconstruct the volumetric internal structure of the cured fibers, generating orthogonal projections (xz- and yz-planes) that verify the preservation of distinct fluid layers throughout the solid fiber (Figure 5Aii). To quantify the manufacturing consistency along the fiber axis, we discretized the xz-plane cross-section into a 3 *×* 2 grid of regions of interest (ROIs) and tracked the mean fluorescence intensity of each sector over the fiber length (Figure 5Aiii). The resulting profiles demonstrate relatively stable signals; for instance, the targeted actuation zone (R1C2) maintained a stable, high-intensity signal, while adjacent non-actuated regions (e.g., R1C3) remained at baseline levels. The observed variation, attributable to minor measurement artifacts rather than structural defects, confirms that ActiSculpt maintains sculpted cross-section fidelity through the liquid-to-solid transition.

**Figure 5:**
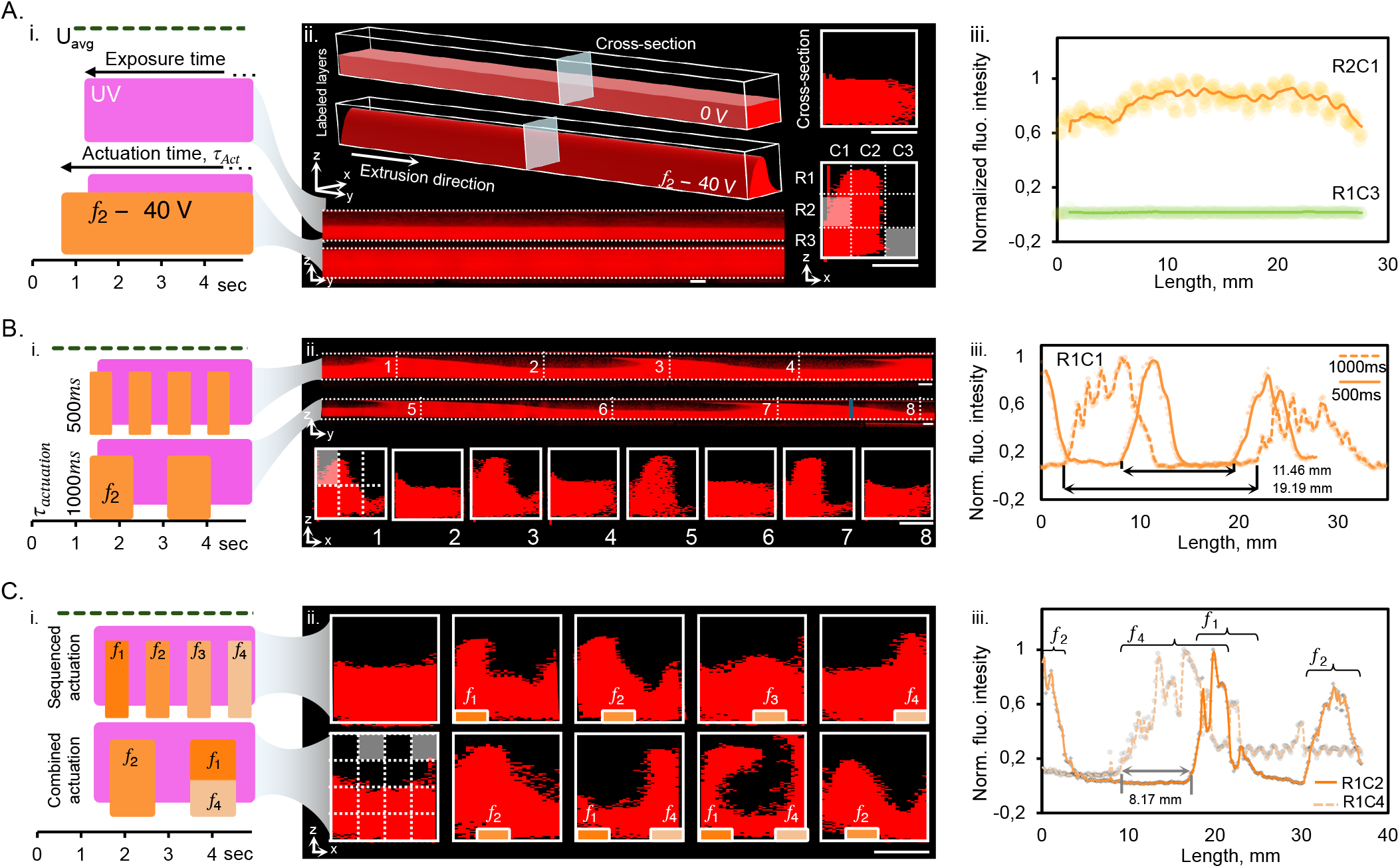
Fabrication of fibrous structures with spatiotemporal actuation. A. Confocal microscopy characterization of fiber cross-sections fabricated with and without acoustic actuation. i. Time sequence of the UV light and acoustic actuation for fiber extrusion. ii. Three-dimensional representative schematic and yz- and xz-plane cross-sectional confocal reconstructions *via* labeled polymer layer tracking; the xz-plane image is segmented into nine regions (3 *×* 3) to quantify pattern stability. iii. Normalized intensity of the selected regions in the continuously actuated extrusion, *f*_2_ at 40 V, demonstrating consistent signal levels along the fiber length. B. Temporal response of acoustic sculpting under cyclic actuation. i. The acoustic actuation was modulated in 500 ms and 1000 ms intervals, producing dynamically sculpted cross-sections. ii. Confocal slices at different positions reveal ON and OFF states across actuation cycles. iii. The average normalized intensity from a selected region (R1C1) displays fluctuation periods (11.47 *±* 0.25 mm and 19.20 *±* 0.57 mm, *n* = 3) aligned with the expected time-of-flight-derived spacing (analytical values of 10 and 20 mm), confirming the deterministic nature of the response. C. Spatial programming of fiber shape *via* sequential and combinatorial transducer actuation. i. Dynamic actuation schemes implemented either by sweeping through excitation locations *f*_1−4_ or by alternating between *f*_2_ and *f*_1,4_. ii. Spatially coded confocal cross-sections capture localized displacements. iii. Cross-sections segmented into 16 regions (4 *×* 4) to track spatial shifts; time-lagged responses between *f*_1_ and *f*_4_ were quantified using regions R1C2 and R1C4. The measured delay (~8.17 *±* 0.24 mm, *n* = 3) aligned with theoretical predictions (7.85 mm), validating spatial control fidelity. Scale bars: 500 µm.

### Dynamic encoding of longitudinal morphology

Beyond static extrusion, ActiSculpt enables the dynamic modulation of fiber topology by temporally varying the acoustic drive. To demonstrate this, we implemented a pulsed actuation protocol (*f*_2_, on-off cycles of 500, 1000, and 2000 ms), effectively encoding a wave deformation pattern along the fiber’s longitudinal axis (Figure 5Bi). Confocal reconstruction of the resulting filaments reveals distinct periodic transitions between the sculpted (actuated) and reference (relaxed) cross-sections (see Movie S1). By extracting cross-sectional profiles at eight discrete locations (Lines 1-8), we confirmed the sharp structural contrast between the alternating states (Figure 5Bii). To quantify the temporal fidelity of this encoding, we correlated the spatial periodicity of the intensity peaks with the theoretical time-of-flight predictions. For an average flow velocity of 9.47 mm*/*s (*Q* = 600 µL min^−1^, ~1038 *×* 1017 µm^2^ cross section), the imposed modulation pulses of 500 and 1000 ms, also considering the non-actuated durations and total signal lag of 0.25 s, yielded spatial length periods of 11.84 mm (measured 11.47 *±* 0.25 mm) and 21.78 mm (measured 19.20 *±* 0.57 mm), respectively (Figure 5Biii). This sub-ten-percent agreement confirms that the fluidic inertia is low enough to map temporal acoustic sequences into precise spatial material gradients. The residual error is attributable to cross-sectional variation from partial clogging during extrusion. An extended characterization of this pulse-shaping behavior is provided in the Supplementary Information (Figures S10-S11, Note S11).

### Spatiotemporal programming of fiber anisotropy

To demonstrate full spatiotemporal control over the fiber architecture, we integrated the temporal modulation capabilities with the system’s spatial addressability. We executed a sequenced actuation that swept the acoustic focus across all four transducer locations (*f*_1_ → *f*_4_) in 500 ms pulses, followed by a combinatorial mode alternating between central (*f*_2_) and outer (*f*_1,4_) IDTs (Figure 5Ci). Confocal analysis of the resulting fibers reveals a shape-coded’ continuum where the cross-sectional geometry evolves deterministically along the extrusion axis. To quantify these transitions, we discretized the cross-section into a 4 *×* 4 intensity matrix (Figure 5Cii) and tracked intensity shifts in specific quadrants (e.g., R1C2 for *f*_1,2_ vs. R1C4 for *f*_4_). The analysis uncovered a distinct spatial phase delay between features generated by upstream (*f*_4_) and downstream (*f*_1_) transducers, a consequence of the finite transit time of the polymer through the actuation zones. The experimentally measured lag distance of ~ 8.17 *±* 0.244 mm aligned closely with the theoretical prediction of 7.85 mm (Figure 5Ciii, Note S9). This predictable offset establishes that ActiSculpt can encode complex, multi-material motifs into continuous fibers with precise longitudinal registration, supporting applications from functional textiles with spatially programmed mechanics to anti-counterfeiting filaments with embedded morphological signatures.

### Translation into a functional 3D printer

To translate acoustic sculpting from bench-scale characterization into an operating additive manufacturing process, we redesigned the platform’s control electronics for integration onto a motion-controlled 3D printer, replacing benchtop instrumentation with a compact, PCB-based architecture. The commercial vector network analyzer was substituted with an inline reflection-coefficient (*S*_11_) monitoring module comprising a bi-directional coupler and power meters, and the collimated UV light source was replaced with a custom PCB-mounted LED array, together reducing the footprint of the acoustic and optical control hardware sufficiently for direct mounting on a printer extrusion head (Figure 6A). The resulting ActiSculpt nozzle was assembled onto a modified 3D printer; two independent, rear-mounted syringe pumps delivered curable and non-curable precursor streams to the nozzle inlet, a microcontroller coordinated acoustic actuation with extruder motion, and an in-line digital microscope mounted on the printer’s movable arm provided flexible real-time imaging of the printed output (Figure 6B; Movie S3).

**Figure 6:**
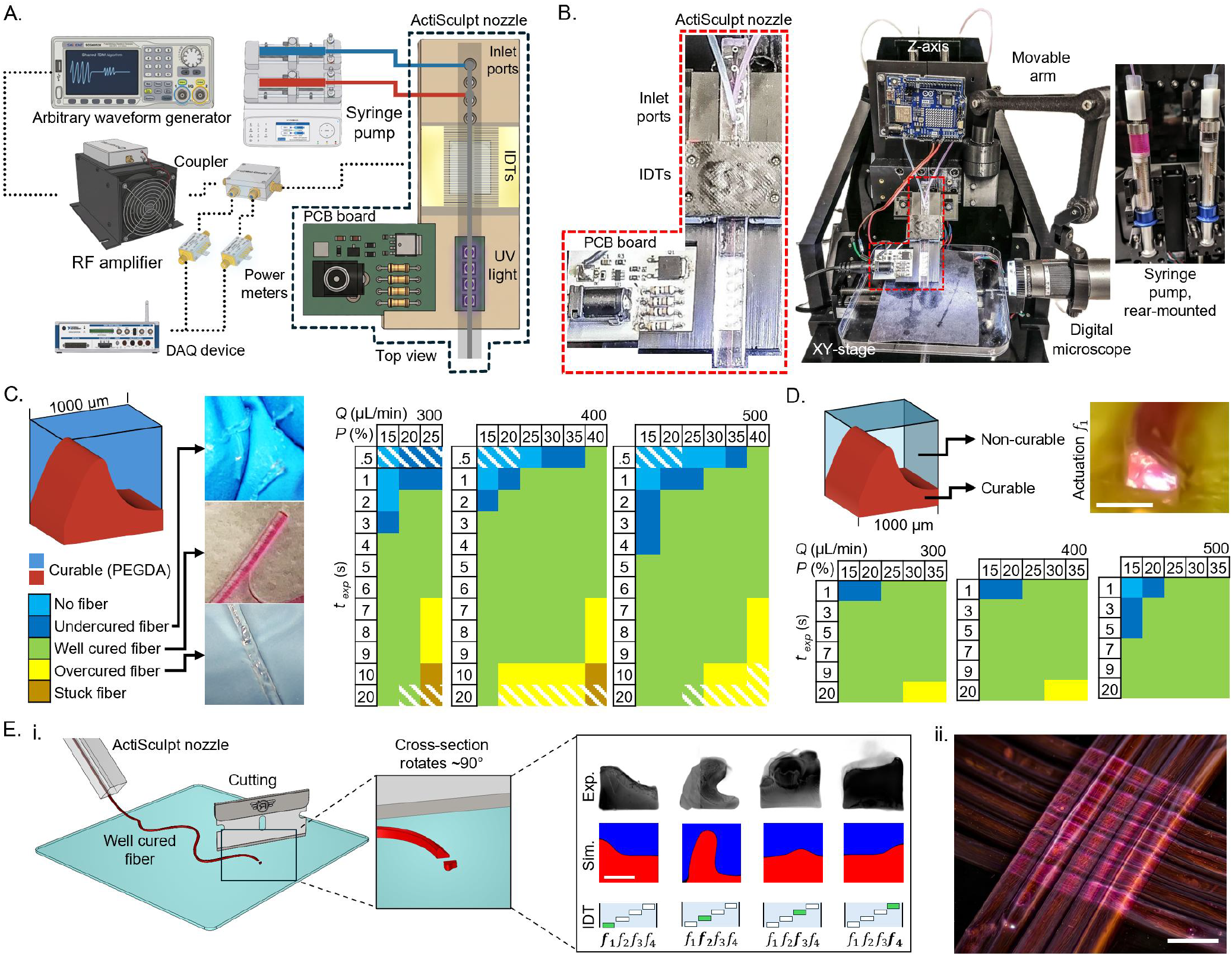
Translation of acoustic flow sculpting into a functional 3D-printing platform. A. Miniaturized footprint of active flow lithography: a custom PCB-based UV LED driver and an inline *S*_11_ monitoring module (bi-directional coupler, power meters) replace benchtop instrumentation, enabling integration of the arbitrary waveform generator, RF amplifier, and DAQ device with the nozzle chip. B. The assembled 3D-printing system: rear-mounted syringe pumps for curable and non-curable precursor streams, the ActiSculpt nozzle mounted on a Z-axis stage for vertical motion, with an XY stage providing horizontal motion to achieve overall three-dimensional positioning, and an in-line digital microscope mounted on the printer’s movable arm, providing the flexibility for real-time imaging of the printed output. C. Printability optimization of a fully cured cross-section: printability matrix for PEGDA fibers across flow rate (*Q*, 300 µL min^−1^ to 500 µL min^−1^), exposure time (*t*_*exp*_, s), and UV power (*P*, %), classifying outcomes as no fiber, undercured, well cured, overcured, or stuck. D. Printability of a half-curable layer configuration: corresponding printability matrix for a co-flowed curable/non-curable stream, showing an expanded well-cured operating window relative to (C); inset shows actuation of transducer *f*_1_ (scale bar: 1 mm). E. Fabricated half-curable fiber cross-sections under individual actuation of each IDT. (i) Schematic of the cross-sectional sampling method: printed fibers were cut perpendicular to their length, rotating the viewing plane ~ 90° to expose the internal cross-section; experimental cross-sections (top row) correspond to single-transducer actuation at *f*_1_ = 37.45 MHz (200 mV pre-amplification), *f*_2_ = 40.645 MHz, *f*_3_ = 44.531 MHz, and *f*_4_ = 48.952 MHz (250 mV pre-amplification for *f*_2_-*f*_4_), at *Q* = 300 µL min^−1^ in all cases, with corresponding simulated cross-sectional profiles (middle row) and the active transducer indicated below each panel (bottom row). (ii) Printed pattern comprising two stacked fiber layers, demonstrating structural assembly of the sculpted fibers into a multi-layer construct. Scale bars: 500 µm (i), 2 mm (ii).

To establish reliable printer operating conditions, we first mapped the printability of a fully cured, single-material (PEGDA) cross-section across *Q* (300, 400, and 500 µL min^−1^), *t*_*exp*_, and UV power (*P*), classifying each combination as yielding no fiber, an undercured fiber, a well-cured fiber, an over-cured fiber, or a stuck fiber (Figure 6C). The well-cured regime occupied a broad central band across all three *Q* values; however, the undercured region (corresponding to insufficient exposure at the shorter timescales tested) extended to longer *t*_*exp*_ as *Q* increased from 300 to 500 µL min^−1^, consistent with the reduced residence time available for photopolymerization at higher throughput. At the longest *t*_*exp*_ (10 and 20 s) and lowest *P*, fibers additionally became overcured or, in isolated cases, stuck within the nozzle.

We then extended this printability characterization to a half-curable, two-stream configuration, in which a curable PEGDA layer was co-flowed with a non-curable stream (Figure 6D). Relative to the fully cured, single-material case, the half-curable configuration exhibited a substantially narrower under-cured region, confined to the shortest *t*_*exp*_ (1 s) at low *P* across all three *Q* values, and no instances of stuck fibers were observed within the tested parameter space. This indicates that the reduced curable-material volume in the half-curable configuration relaxes the *t*_*exp*_ and *P* constraints required to achieve a well-cured fiber, broadening the accessible printing window relative to the fully cured case.

Having established robust printability of the half-curable configuration, we assessed whether the cross-sectional sculpting demonstrated on-chip (Figures 1-5) is retained once translated into the printed fiber output, sampled via the cross-sectioning method shown in Figure 6Ei. Actuating each of the four IDTs individually while printing the half-curable stream produced four distinct, IDT-specific fiber cross-sectional geometries, and corresponding numerical simulations of the sculpted interface closely reproduced each experimentally observed cross-section (Figure 6Ei), confirming that acoustic streaming remains the dominant sculpting mechanism once translated from bench-scale characterization into the printed fiber. Finally, to demonstrate that this printed, acoustically sculpted output extends beyond a single filament, we assembled multiple printed fibers into a two-layer stacked structure (Figure 6Eii), confirming that active sculpting does not compromise the fiber’s capacity for structural assembly into larger, multi-fiber constructs.

## Discussion

In summary, we have introduced ActiSculpt, a lithographic platform that replaces flow-shaping static physical structures (e.g., pillars, nozzles, or herring-bone patterns) with dynamic acoustic-sculpting operators. By integrating tunable acoustic streaming with stop-flow and continuous lithography, the system decouples fluid deformation from chaotic mixing, enabling the deterministic sculpting of cross-sectional architectures. This approach overcomes limitations of traditional passive flow shaping strategies, offering a new method without issues of high pressure drop or the constraints of one-design-one-function devices, while enabling real-time programmability.

We have demonstrated sculpting capability across two domains: steady-state cross-sectional shape design and dynamic, real-time control. In the steady-state domain, material geometric properties, such as moment of inertia, can be precisely tuned, enabling potential control over the mechanical stiffness of microstructures. In the dynamic domain, acoustic programmability enables the manufacture of different particle or fiber shapes from the same device —a one-device, many-function capability unavailable in passive systems. Temporal modulation of the acoustic field extends this further, enabling spatiotemporal sculpting that encodes complex morphology along the fiber axis, yielding longitudinally patterned filaments that are beyond the reach of conventional extrusion methods.

Beyond bench-scale characterization, we further demonstrate that this sculpting capability translates directly into an operating additive manufacturing process. Integrating the ActiSculpt nozzle onto a motion-controlled 3D printer, we established printability windows for both single-material and half-curable, two-stream fiber configurations, and confirmed that acoustically programmed cross-sections, together with their capacity for assembly into multi-fiber structures, are preserved once printed. This translation moves the platform beyond bench-scale, chip-based characterization toward a functional additive manufacturing process, directly addressing a limitation shared by existing acoustofluidic and passive flow-sculpting techniques, which have so far been demonstrated only at the bench.

These capabilities position ActiSculpt as a flexible platform for functional material manufacturing, extending flow lithography into a domain of active, real-time morphological control. The ability to embed spatially encoded morphological information and functional anisotropy directly into the fiber core opens new avenues for smart textiles, where mechanical or optical properties vary along the weave, and anti-counterfeiting applications, where unique morphological signatures serve as intrinsic security keys.

## Materials and Methods

### Fabrication of the microchannel

Microfluidic channels featuring a 1 *×* 1 mm^2^ square cross-section and configurable multi-inlet architectures were fabricated via soft lithography using a hybrid mold assembly. The master channel features were patterned using a high-resolution resin 3D printer (Sonic Mini 8K, Phrozen). To prevent PDMS curing inhibition caused by residual photoinitiators, the resin molds underwent a strict post-processing protocol involving UV exposure, thermal annealing, and surface passivation, as described by Venzac et al. [59]. The mold housing was fabricated from acrylonitrile butadiene styrene (ABS) using fused deposition modeling (X1 Carbon, Bambu Lab) to ensure thermal stability. Polydimethylsiloxane (PDMS) was mixed at a 1:10 base-to-agent ratio, cast into the mold, and cured at 65 °C for one hour. The device was sealed against a flexible 150 µm-thick PDMS membrane, prepared by spin-coating the prepolymer onto a silanized silicon wafer (500 rpm, 30 s), via oxygen plasma bonding (500 W, 60 s, 0.6 Torr). This membrane thickness is critically tuned to ensure acoustic transparency and mechanical stability, avoiding the high-damping regime observed at thickness-to-wavelength ratios above ~ *O* (10) [60].

#### Fabrication of interdigital transducers

Aluminum interdigital transducers were patterned on lithium niobate (LiNbO_3_, 128°YX, 31, 0.5 mm thickness, Hangzhou Freqcontrol Electronic Technology Ltd.) piezoelectric substrates (converting the electric input into acoustic waves by the inverse-piezoelectric effect) via a subtractive photolithography process [61–63]. First, the substrates were cleaned with acetone and isopropanol, followed by the uniform deposition of an aluminum thin film using magnetron sputtering (NanoPVD, Moorfield Nanotechnology). To pattern the electrodes, the samples were dehydrated (120 °C), spin-coated with AZ5214E positive photoresist (5000 rpm, ~1.25 µm thickness), and soft-baked at 100 °C for 3 min. Lithographic exposure was performed using a maskless laser writer (*µ*MLA, Heidelberg Instruments), followed by development in AZ351B (1:4 dilution). Prior to wet etching, the photoresist mask was stabilized via a hard bake at 130 °C for 4 min. Finally, the specific electrode geometries were defined by wet chemical etching of the exposed aluminum, followed by resist stripping and rinsing. Intended wavelengths for each IDT were 104, 96, 88, and 80 µm, corresponding to ideal resonant frequencies of 38.27, 41.45, 45.23, and 49.75 MHz, respectively, under the 3980 m*/*s speed of sound in the LiNbO_3_ substrate provided by the manufacturer. Since manufactured geometries were slightly different from the intended values, resonant frequencies were measured v a vector network analyzer as 37.32, 40.32, 44.07, and 48.42 MHz, respectively.

### Device assembly and integration

To ensure precise alignment and reliable electrical interfacing between the IDT substrate and the PDMS microchannel, a custom modular holder was designed and fabricated. The assembly features a recessed base for substrate registration and a dedicated slot for inserting 3D-printed physical photomasks. Electrical connection to the transducer pads is established via a discrete pogo-pin array, which is secured against the substrate using a sandwich-style clamping lid to ensure uniform mechanical pressure and acoustic coupling. The structural components were fabricated using fused deposition modeling (PLA) for the primary chassis, while high-resolution stereolithography (resin) was employed for fine alignment features requiring tighter tolerances. A detailed schematic of the full device assembly is provided in Figure S1.

### Experimental setup and instrumentation

The platform architecture synchronizes acoustic actuation, fluid drive, and photopolymerization via a centralized control interface. Acoustic waveforms were synthesized by an arbitrary function generator (SDG6022X, Siglent), amplified (LZY-22+, Mini-Circuits), and routed to the transducers through a 4-port mechanical relay. This switching topology, governed by a data acquisition unit (USB6003, National Instruments), enabled dynamic transitions between high-power actuation and real-time spectral characterization via a vector network analyzer (SVA1000X, Siglent). Fluid injection was regulated by a precision syringe pump (Fusion 4000, Chemyx), while photopolymerization was triggered by a collimated UV LED source (AC475, OmniCure) for a defined UV exposure time (*t*_*exp*_, in s). A custom LabVIEV environment, augmented with embedded Python algorithms, orchestrated the system timing. This master controller managed the time-division multiplexing (TDM) sequences and synchronized the specific operational modes: executing synchronized flow-stop/UV-flash cycles for particle lithography, and maintaining continuous irradiation with concurrent acoustic drive for fiber extrusion. To standardize the propagation across different experimental conditions, the TDM algorithm synthesizes arbitrary waveforms based on the signal frequency (*f*), amplitude (*V*), and sampling rate (*f*_*s*_). The signal structure was set to a fixed burst of *N* = 30 cycles to maintain a constant number of wave-propagation cycles per actuation event. Consequently, the algorithm dynamically adjusts the sampling rate to lock the duty cycle of each active channel at ≈ 30%, providing similar resting time for heat removal, governed by the relationship:

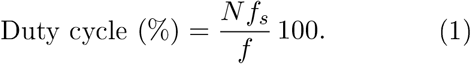

### Precursor solution preparation

The photopolymerizable ink consisted of polyethylene glycol diacrylate (PEGDA, *M*_*n*_ ≈ 575) dissolved in deionized water (1:1 v/v). To enable multicolor fluorescence tracking, separate precursor aliquots were doped with methacryloxyethyl thiocarbamoyl rhodamine B (0.003% w/v) for red emission, and a custom methacrylate-functionalized dye, MA-P4VB (2-(methacryloyloxy)propyl 6-(4-methoxy-2,5-bis((E)-2-(pyridin-4-yl)vinyl)phenoxy)hexanoate, 0.006% w/v), for blue emission [64]. Lithium phenyl-2,4,6-trimethylbenzoylphosphinate (LAP) was added to all solutions at a concentration of 0.1% w/v to serve as the water-soluble photoinitiator.

## Acknowledgments

This work is funded by the Deutsche Forschungsgemeinschaft (DFG, German Research Foundation) - Project number 567108437 and the National Research Foundation of Korea (NRF) grant funded by the Korea government (MSIT) (No. RS-2025-02316565).

## Author Contributions

G.D. and M.A.S. conceived the study and designed the platform architecture. M.A.S. designed and fabricated the microfluidic devices and interdigital transducers. M.A.S., S.H., A.C., and L.E. performed all sculpting, particle, and fiber-fabrication experiments, developed the image-analysis pipelines, and analyzed the data. M.A.S., D.S., and G.D. designed and validated the numerical model of the acoustic-streaming flow field and contributed to the interpretation of the cross-sectional sculpting results. J.P. and G.D. contributed to the acoustofluidic design and resonator characterization and secured funding for this work. G.D. supervised the project. M.A.S., S.H., and G.D. wrote the manuscript with input from all authors. All authors discussed the results and approved the final version.

## Data Availability

The main data supporting the findings of this study are available within the paper and its Supplementary Information. Other datasets generated or analyzed during the current study are available from the corresponding author on request. Source data are provided with this paper.

## Code Availability

The custom Python-based image-analysis pipelines used in this study-the particle cross-section analysis pipeline (https://github.com/masahinphd/actisculpt_particleanalysis) and the confocal-stack reslicing pipeline (https://github.com/masahinphd/actisculpt_confocal), and the sculpting characterization (https://github.com/masahinphd/actisculpt_characterization)-are publicly available on GitHub. The numerical model implementation COMSOL Multiphysics and corresponding input files are available from the corresponding author upon reasonable request.

## Supporting Information

**Figure S1:**
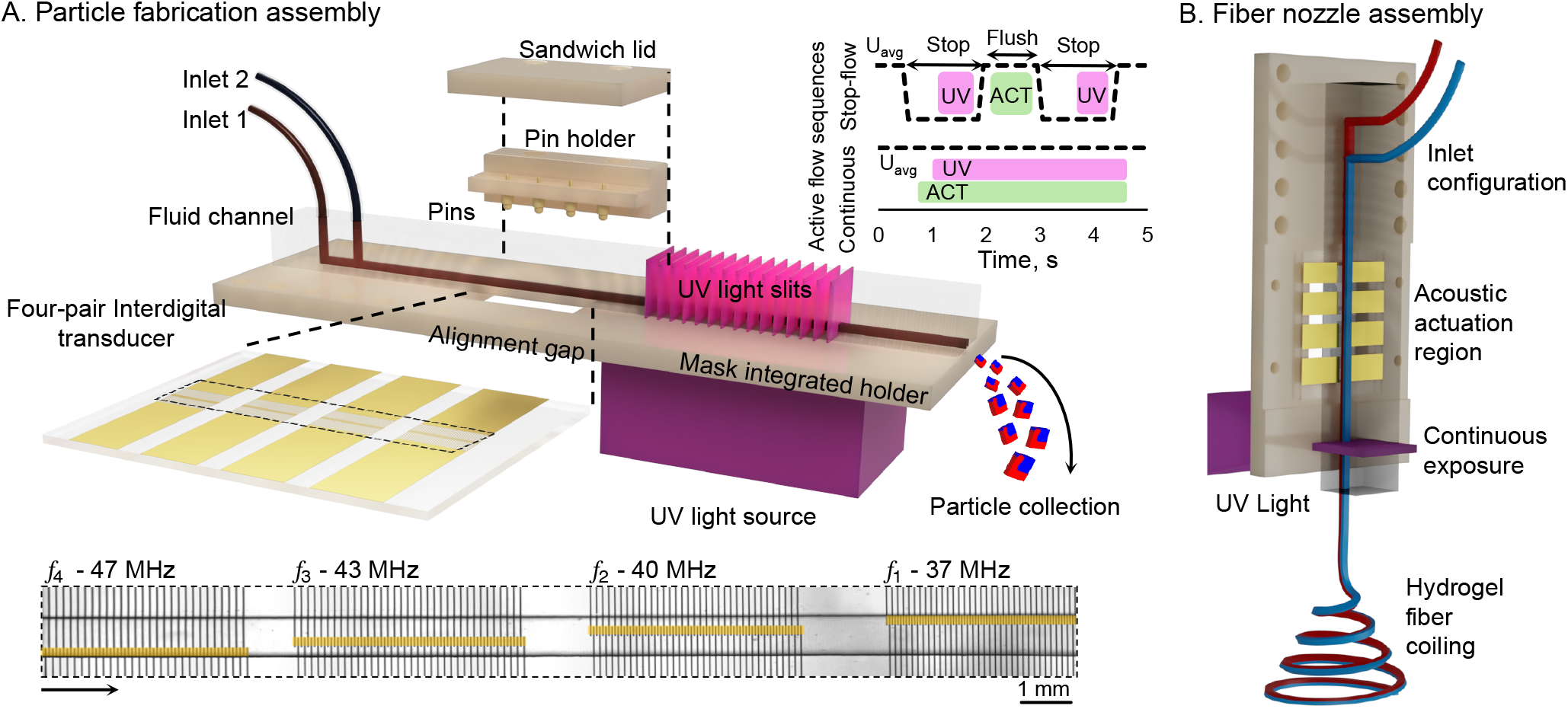
Assembly configuration for particle and fiber fabrication. A. Schematic for the assembled ActiSculpt device, where a sandwich lid compresses the fluid channel towards the four interdigital transducers, avoiding any sliding during the experimental work. A mask-integrated holder allows the alignment of the IDT and PDMS channel via an alignment gap. Stop-flow lithography can be performed to produce particle slices via UV light slits dictated by the mask integrated into the holder. B. Schematic for the fiber nozzle assembly, where continuous UV light is exposed through a single slit after the acoustic actuation region.

## Note S1: Temperature measurements

To enable temperature measurements within the microchannel during acoustic actuation, the microfluidic channel design was modified to include an additional outlet port located downstream of the acoustic interaction region. This redesign facilitated improved flow control and thermal profiling across the region of interest. Two K-type thermocouples were integrated into the system, one upstream and the other down-stream of the acoustic actuation zone, enabling direct measurement of the temperature differential induced by acoustic energy. The thermocouples were connected to a digital thermocouple reader, enabling real-time monitoring of localized thermal changes within the microchannel during operation (Figure S2).

Fluorescently labeled fluid streams were utilized to enable high-contrast imaging using conventional fluorescence microscopy. Following fabrication and washing steps, the microparticles, engineered with an approximate height-to-thickness aspect ratio of 1:4, were collected and seeded into a well plate containing deionized (DI) water supplemented with 5o% w/v Pluronic F127. This medium helped prevent particle aggregation and adhesion to the substrate during imaging. Dual-channel fluorescence detection was employed to separately capture red and blue emission signals corresponding to the incorporated fluorescent dyes. Optical and illumination parameters were carefully optimized to maximize the signal-to-noise ratio, ensuring clear distinction between particle features and background fluorescence. For characterization metrics, particle images were cropped by 10% (~ 20 px to 25 px) from the outline to suppress discrepancies caused by the outer layers. Notably, for the metrics that require stacking or projecting images, i.e., uniformity and cumulative displacement, an unavoidable, small pixel-scale misalignment remains when centering the experimental images.

**Figure S2:**
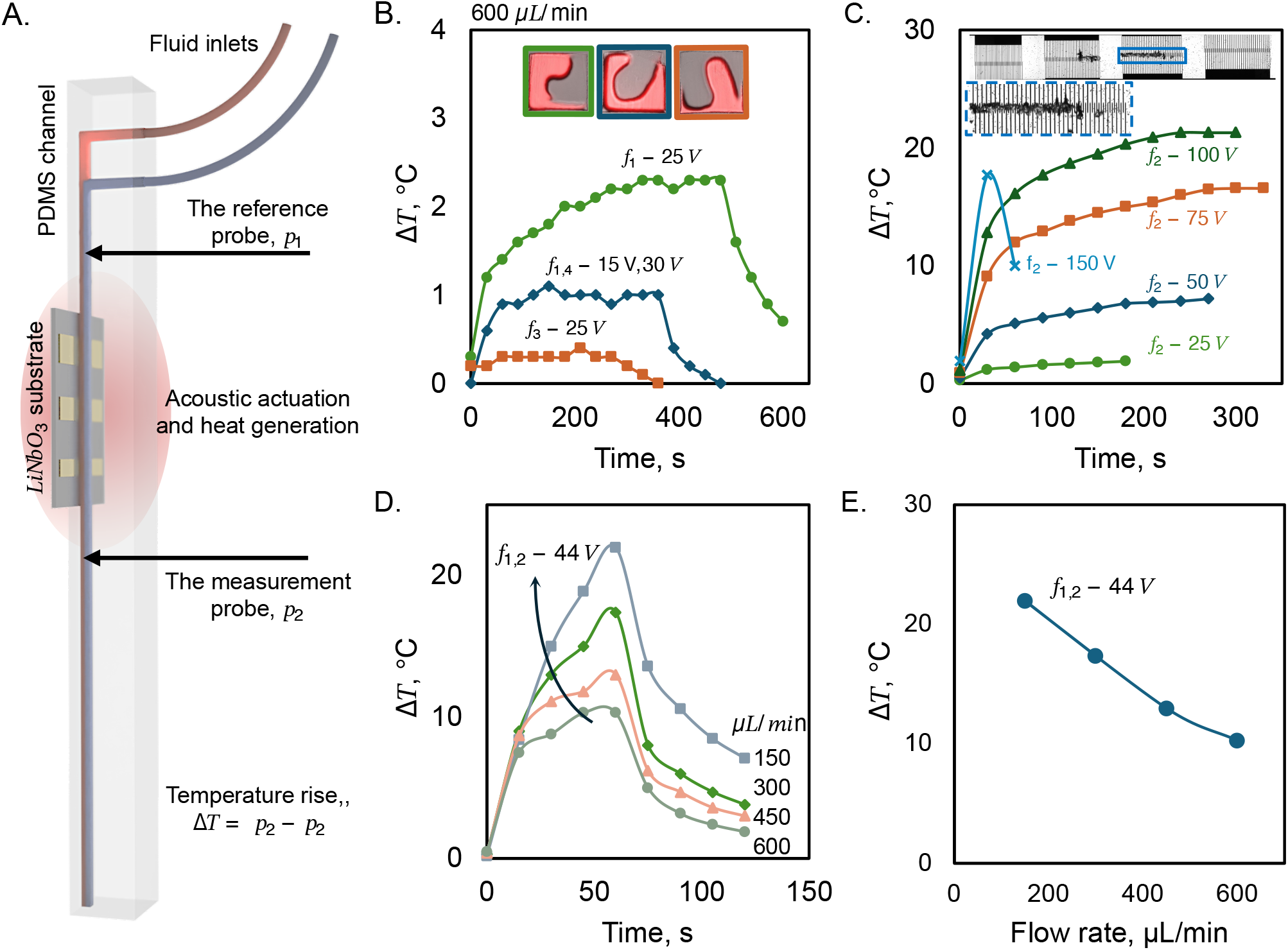
Temperature measurement under different conditions. A. Schematic for the measurement setup for a differential read. B. Long-period measurement with a reasonable amount of voltage input (15 V to 30 V) at *Q* = 600 µL min^−1^, with corresponding images depicted as color-coded insets. C. Durability test under high-voltage application on the selected actuator, *f*_2_, varying input levels from 25 to 150 V for at least 5 minutes; inset displays the defect that occurred when 150 V high actuation was applied. D. Dual actuation of *f*_1,2_ at 44 V at varying *Q*. E. Dual actuation plotted along *Q*.

## Note S2: Imaging the particles

Facilitating the unbounded-medium assumption by using a smaller actuation width (*D*_2_) than the channel width (*D*_1_) enables applying the momentum-force approach to model acoustic streaming as an analytically calculated body force based on the acoustic settings and propagation medium properties. Force approaches have been developed and advanced by many researchers; however, we used Nyborg’s notation and the implementation presented by Mason [S1] for our approach. The force induced by acoustic wave propagation can be written as

**Figure S3:**
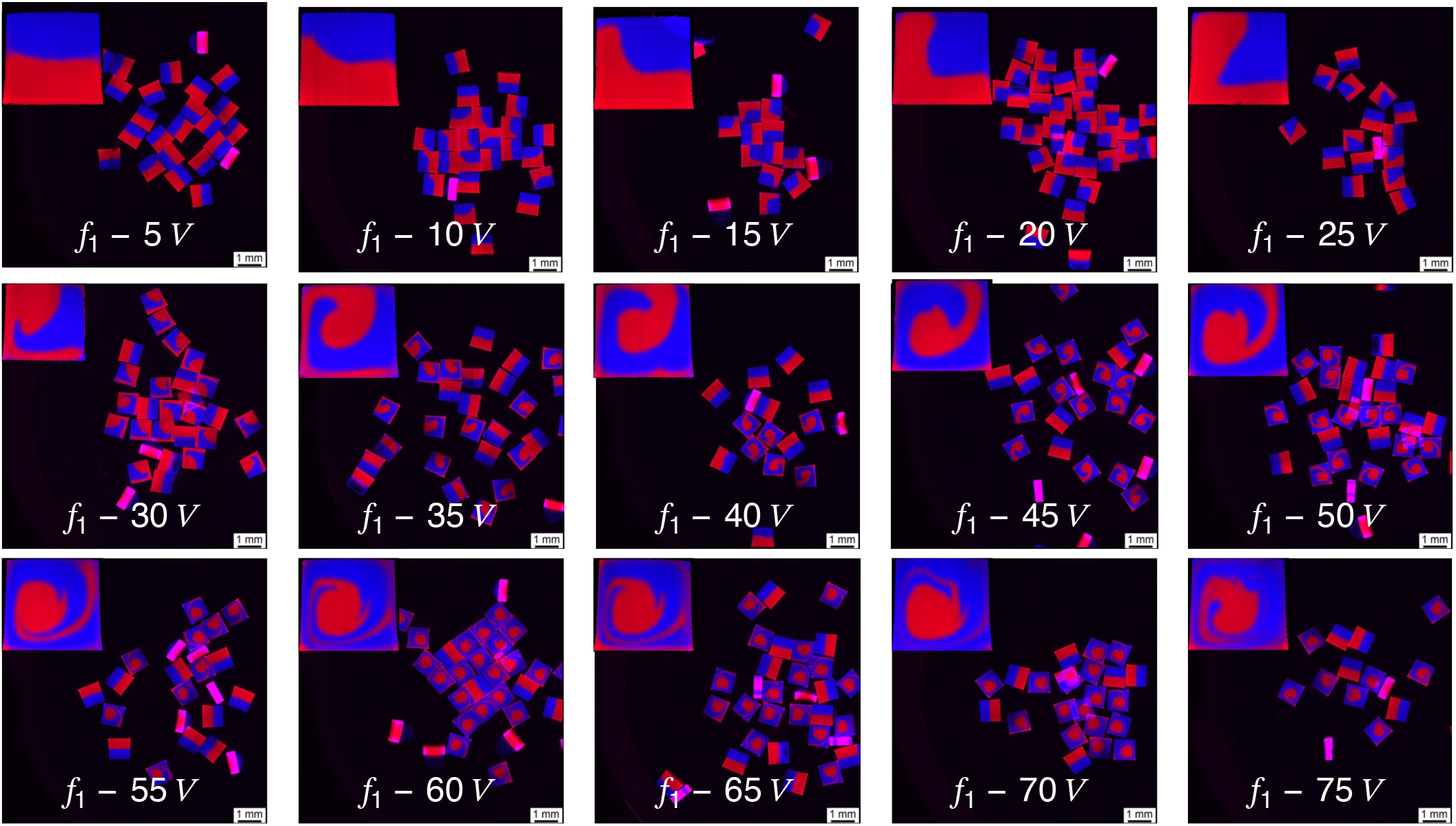
A library for the actuation site of *f*_1_ at varying voltage input values. Fabricated particles are seeded in a well plate for traditional planar fluorescent microscope imaging. A post-processed single-particle example is shown at the top left of each well-plate image. The experiment at 480 mV input power was repeated once more to demonstrate batch-to-batch stability. Cropped images are not to scale; the scale bar on the well-plate images is 1 mm.

**Figure S4:**
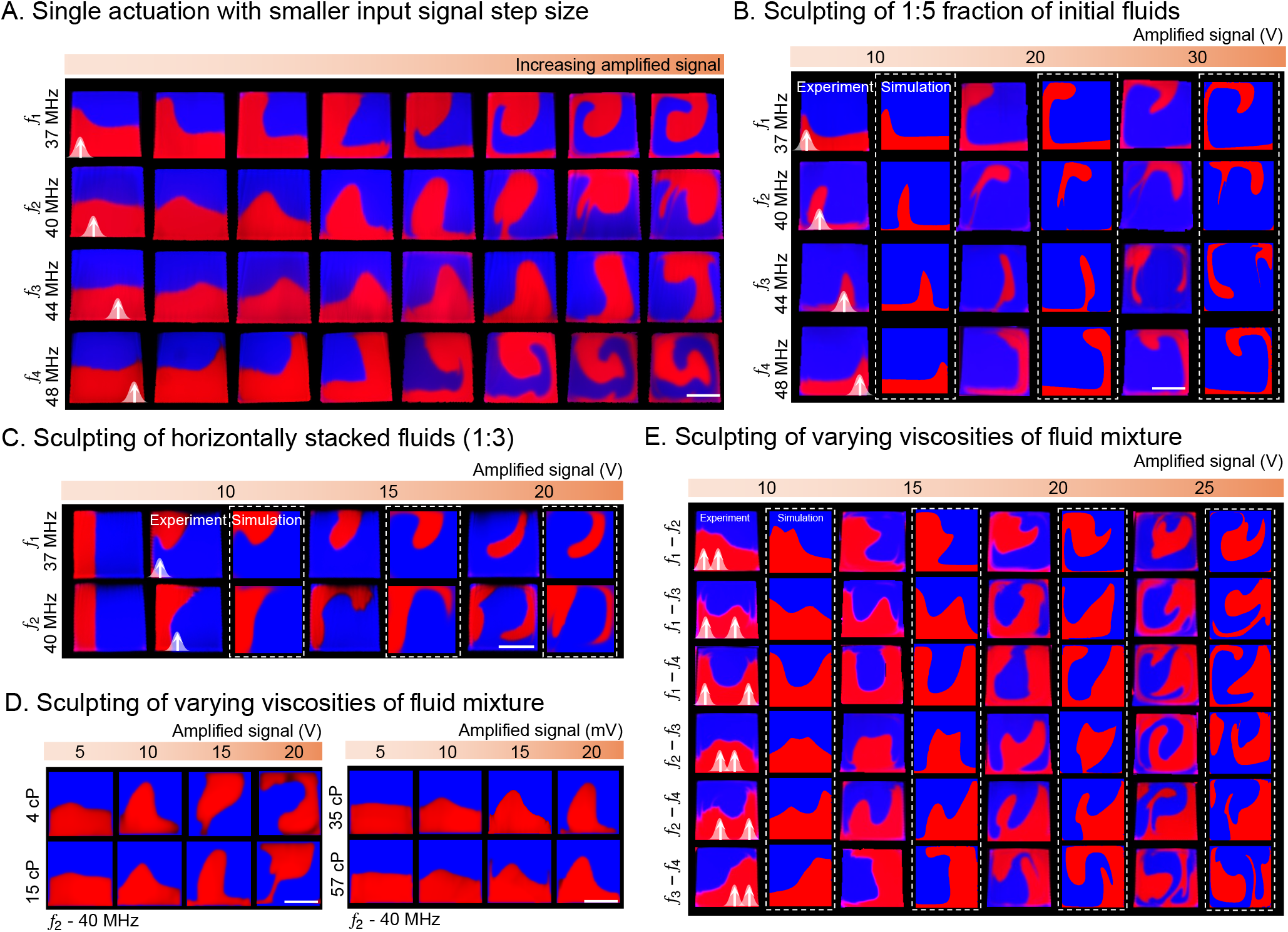
The extent of the generated ActiSculpt library. A. Example of the actuation sweep with increasingly smaller input signal. B. Additional example of the actuation sweep with the fluid fraction dictated by a flow ratio of 1:5. C. Additional example of the actuation sweep with the fluids stacked horizontally and the fluid fraction dictated by a flow ratio of 1:3. D. Experiments conducted with different mixtures of PEGDA and water; estimated viscosities vary from 5 to 50 cP. Only the bottom layer is fluorescently labeled. Scale bar represents 500 µm. E. Library generated by combinatorial dual actuations, sweeping the input voltage from 80 to 240 mV, left to right. Scale bar represents 500 µm.

## Note S3: Building a numerical model

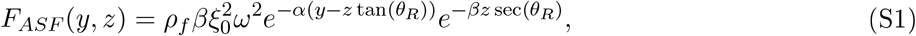

where *ρ*_*f*_ is the density of the fluid, *ξ*_0_ is the initial displacement amplitude, *ω* is the angular frequency, *θ*_*R*_ is the Rayleigh angle, and *α* and *β* are the attenuation coefficients at the interface of the media and in the fluid, respectively. The Rayleigh angle describes the propagation angle of a surface acoustic wave into a fluidic medium, defines the geometric limits of this equation, and is observable experimentally in an anechoic chamber [S2-S4]. The variables and equations were implemented in COMSOL Multiphysics. The ‘Laminar Flow’ module solves the induced Navier-Stokes equation to calculate velocity and pressure distributions, including the acoustic streaming effect. In a simple square cross-sectional channel, four IDT positions are located as in the ActiSculpt device and are modeled as separate domains to define a body-force source.

**Figure S5:**
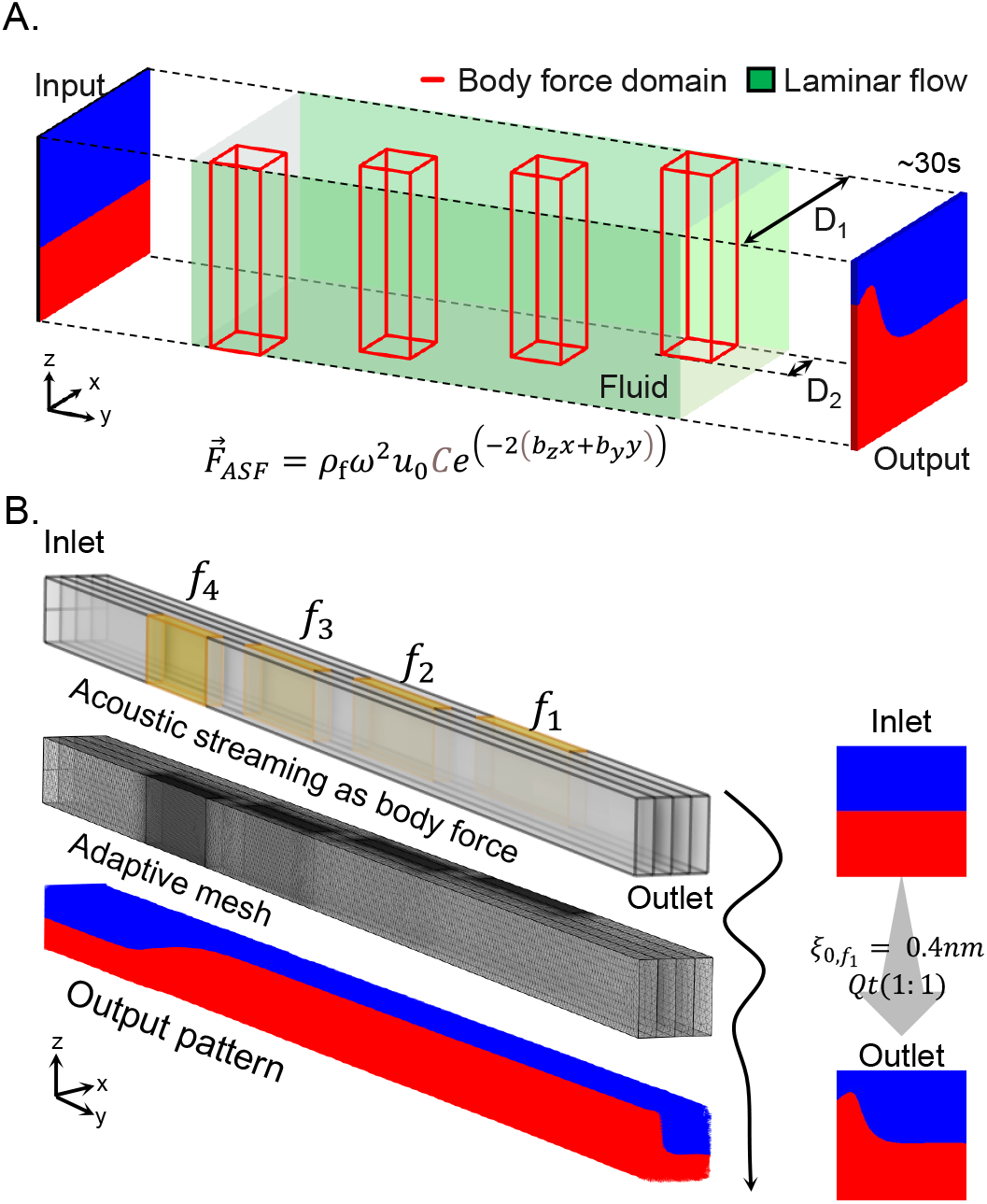
3D numerical modelling of the acoustic streaming. A. Domain creation of the acoustic streaming as a body force in the laminar flow equations. B. Implementation of the body-force approach in COMSOL Multi-physics. An adaptive mesh is used to generate a finer mesh around the actuation domains.

## Note S4: Post-processing of the experimental 2D fluorescent images of fabricated particle slices

To enable the high-throughput quantitative characterization of fabricated microparticles, we developed a custom Python-based image analysis pipeline to isolate and evaluate individual cross-sections from raw multi-channel fluorescence micrographs (8-bit or 16-bit). The automated workflow initiates by sorting raw image data according to experimental parameters, subsequently applying thresholding and watershed segmentation algorithms to delineate individual particle boundaries. Following the size-based filtration of background noise and imaging artifacts, the algorithm extracts morphological descriptors to classify the population into actuated (sculpted) and non-actuated (reference) subsets. Each classified particle is then subjected to spatial normalization, comprising automated cropping and rotational alignment to a standardized coordinate system, before the discrete fluorescence channels are isolated and exported for statistical evaluation (Figure S6). The entire process takes a few seconds per loaded image. To facilitate usability and open-source reproducibility, a graphical user interface (GUI) was integrated into the core processing script using generative AI models (GitHub Copilot Student), and the complete source code is publicly accessible via a dedicated GitHub repository (https://github.com/masahinphd/actisculpt_particleanalysis).

### Calculation of amplified acoustic output voltage

The time-division multiplexing (TDM) algorithm, implemented in Python, synthesizes discrete sinusoidal burst packets based on the user-defined number of actuation sites and their target amplitudes. The digital amplitude inputs are constrained by the arbitrary waveform generator’s 8-bit vertical resolution (Siglent SDG6022X), which maps integer values from −127 to 127. Throughout this study, the generator’s baseline reference voltage was fixed at 500 mV*/*unit, meaning the maximum digital integer of 127 corresponds directly to a 500 mV*/*unit pre-amplification signal. The synthesized signal is subsequently routed through an RF amplifier (LZY-22+, Mini-Circuits) that provides a constant gain of 44 dB, equivalent to a linear voltage multiplier of approximately 158.49. Consequently, the conversion factor from the user-defined digital unit to the final amplified output voltage (*V*_*out*_) delivered to the transducer is calculated as

**Figure S6:**
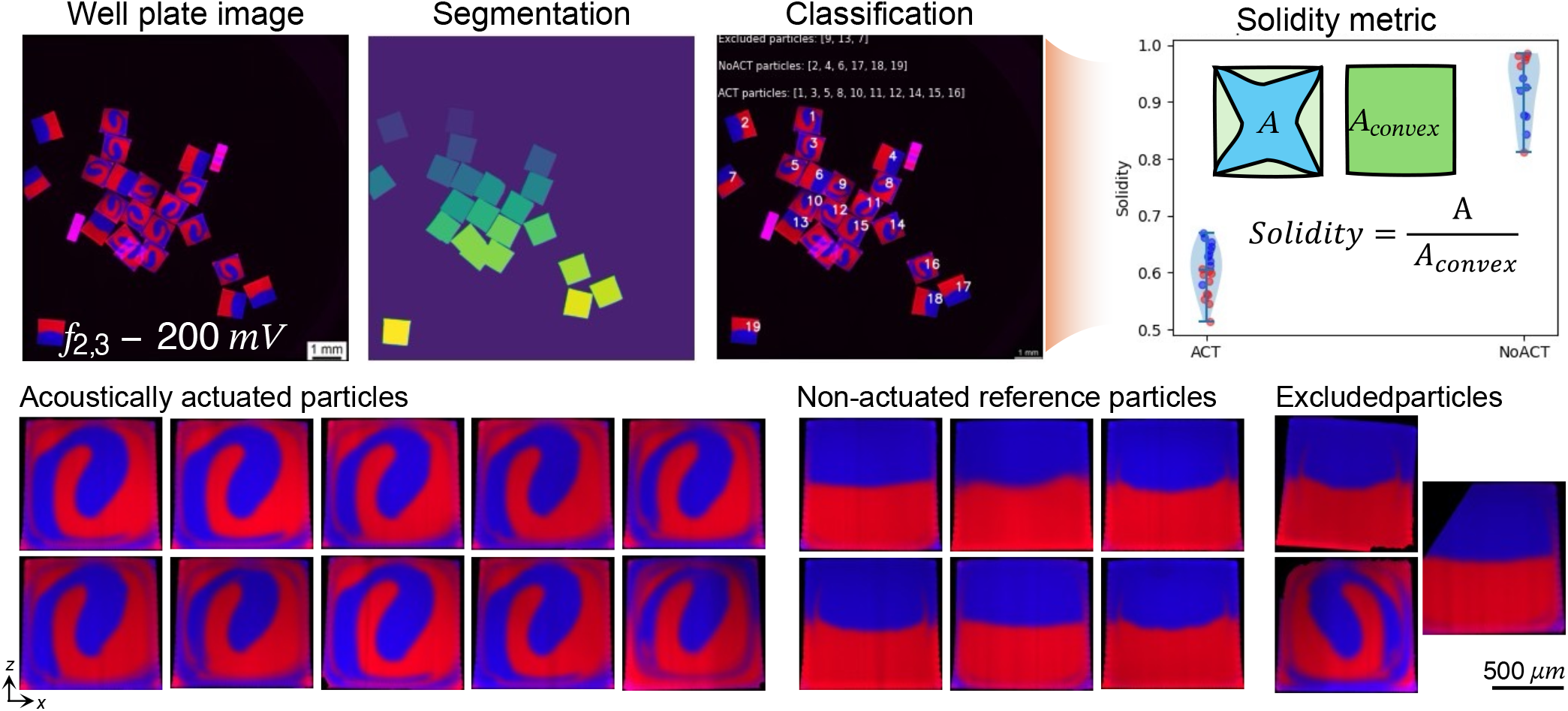
Post-processed particles extracted from the well-plate image of the experiment *f*_2,3_-200 mV. Actuated and reference non-actuated particles are segmented, spatially normalized, and classified depending on the pattern characteristics (circularity, solidity, etc.). Discrepancies present in the particles due to experimental error were also filtered out. Scale bar represents 500 µm.

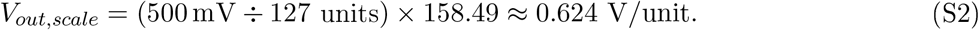

To illustrate, requesting a digital amplitude of 10 units yields an initial generator output of 39.37 mV. After passing through the 44 dB amplifier, the final acoustic driving voltage scales deterministically to 6.24 V (10 units *×* 0.624 V*/*unit).

## Note S5: Mixing and flow regime analysis

To evaluate the extent of fluid mixing versus deterministic sculpting, the cross-sectional fluorescence intensity was analyzed for each isolated particle. Following spatial segmentation, the discrete color channels were separated, and the normalized standard deviation (*σ*) of the pixel intensities was calculated across the entire particle area. In this framework, an ideally separated, binary fluid interface yields *σ* = 0.5, whereas a completely homogenized (perfectly mixed) stream yields *σ* = 0. Throughout the mixing experiments, *Q* was maintained at 600 µL min^−1^, with a 1:1 volumetric flow ratio between the two inlets to ensure consistent kinematic baseline conditions. To characterize the physical flow regime and evaluate the relative dominance of advective versus diffusive transport, we calculated the dimensionless Reynolds (*Re*) and P clet (*Pe*) numbers. The fluid properties were defined as follows: mixture density, *ρ* (1060 kg*/*m^3^); flow velocity, *v*, 0.01 m*/*s; characteristic length, *L*, 0.001 m; mixture viscosity, *µ*, 0.01 Pa s. The Reynolds number, which defines the ratio of inertial to viscous forces, was calculated as *Re* = *ρvL/µ* = 1.06. This low *Re* value (*Re* ≪ 2000) confirms that the system operates firmly within a predictable, laminar flow regime, devoid of turbulent inertial mixing. Furthermore, to quantify the timescale of advection relative to diffusion, the P clet number was evaluated as *Pe* = *vL/D*, where *D* is the diffusion coefficient. The theoretical diffusion coefficient for the fluorescent solute (rhodamine B methacrylate) was estimated using the Stokes-Einstein relation, *D* = *k*_*B*_*T/*(6*πµr*_*h*_). Given the Boltzmann constant (*k*_*B*_ = 1.38 *×* 10^−23^ J*/*K), an absolute temperature (*T*) of 298.15 K, and an estimated hydrodynamic radius (*r*_*h*_) of ~ 1 nm for the dye molecule, this yields a high Péclet number of *Pe* ≈ 4.58 *×* 10^5^. The approximate transverse diffusion length, *x*_*d*_, can also be calculated from 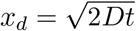, which yields 14.8 µm for a residence time of 5 s (*v* = 0.01 m*/*s, 5 cm channel length). These magnitudes (*Pe* ≫ 1, *x*_*d*_ ≪ *L*) rigorously demonstrate that mass transport within the microchannel is overwhelmingly dominated by advection, ensuring that the sculpted interfaces remain sharply defined and are not degraded by passive molecular diffusion during the transit time.

## Note S6: Batch-to-batch stability and pattern uniformity analysis

To quantify batch-to-batch reproducibility, we established a pixel-wise uniformity metric based on a consensus threshold. Specifically, a spatial pixel was considered part of the conserved structural motif if it was occupied in at least 50% of the analyzed particles within a given batch. This “majority rule” approach was intentionally implemented to mitigate the impact of microscopic registration limits during the automated segmentation and spatial normalization workflows. Because pixel-by-pixel intersection analysis is inherently sensitive to discrete spatial aliasing, even visually imperceptible image translations (e.g., 5-10 pixels) can disproportionately penalize the calculated uniformity score, despite the macroscopic fluid architecture remaining fundamentally stable. By utilizing a consensus threshold, the metric accurately reflects the true morphological stability of the acoustic sculpting process rather than digital alignment artifacts. However, for downstream applications demanding absolute spatial stringency, the accompanying ActiSculpt open-source analysis suite features a “strict uniformity” (100%-pixel match) configuration, enabling users to evaluate image stacks under conditions of absolute pixel-perfect overlap.

## Note S7: Kinematic tracking and fluid displacement analysis

To precisely map the kinematic trajectory of the fluid interface, we conducted high-resolution amplitude (5- or 10-unit → ~3 or ~6 V) sweeps utilizing a smaller voltage step size than the standard 20-unit intervals (~ 12 V). During these granular sweeps, a newly fabricated transducer was integrated for the *f*_4_ actuation band, which exhibited a slight deviation in electromechanical efficiency, as confirmed by *S*_11_ spectral measurements. To normalize the acoustic power across independent experimental batches, we performed comparative image analysis of the resulting cross-sections, establishing an empirical amplitude scaling factor of 1.65 for the *f*_4_ dataset.

When calculating the center-of-mass (CoM) displacement, actuation voltages exceeding 50, 28, 37, and 43 V (for *f*_1_ through *f*_4_, respectively) were intentionally excluded. In these high-power regimes, the deformed fluid interface impinged upon the upper channel wall. This boundary interaction induced complex topological transformations, such as severe phase splitting and spiral vortex formation, that precluded reliable automated segmentation for center-of-mass calculations.

To quantify the spatial resolution of the CoM shifts, linear regression was applied to the displacement data strictly within the unconstrained deformation regime. This linear fit was bounded by the voltage thresholds (*<* 50 V for *f*_1,2,3,4_). Restricting the analysis to these bounds effectively isolates the primary, streaming-driven deformation phase, minimizing nonlinear hydrodynamic disturbances caused by the channel walls’ geometric confinement (Figure S7C).

**Figure S7:**
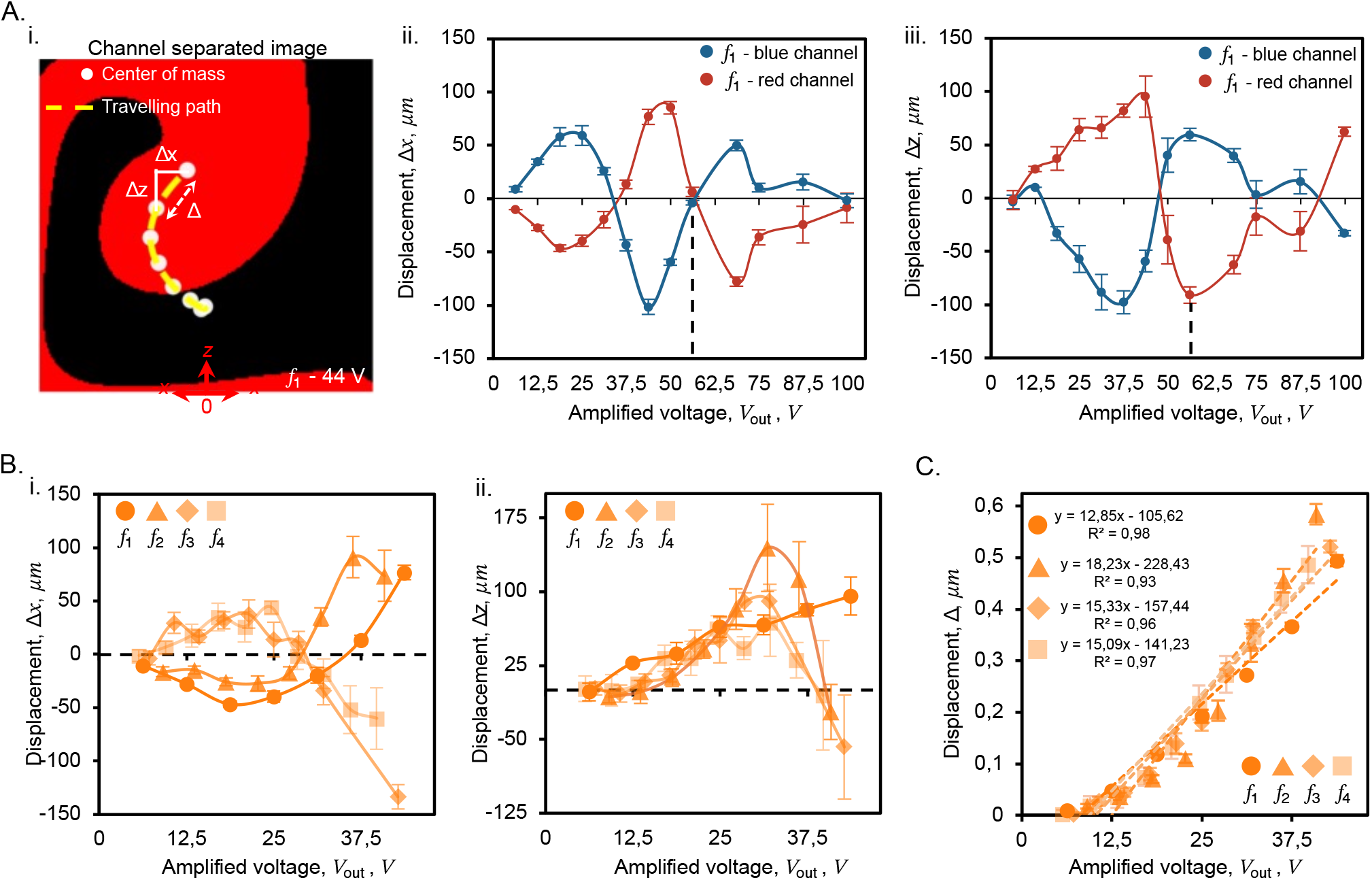
Fluid displacement kinematics. A. The calculation method: locating the center-of-mass points and measuring the distance between each center (i); separated-axis plots for a wider amplitude dataset on *f*_1_ over the x-axis (ii) and z-axis (iii), respectively. B. Representation of all actuation sites, *f*_1_ → *f*_4_, over the x-axis (i) and z-axis (ii). C. Calculation of the displacement per voltage for each actuation site.

## Note S8: Calculation of the second moments of area about the geometric centroid

To evaluate the mechanical implications of the sculpted cross-sections, we calculated the area moments of inertia (*I*_*x*_, *I*_*z*_), which govern a structure’s resistance to bending, and the polar moment of inertia (*I*_*y*_), which dictates resistance to torsion. Isolated particle images were first separated into discrete fluorescence channels and binarized. Because diffusive mixing between the fluid phases was negligible, binarization was efficiently achieved by subtracting the complementary fluorescence channels. To eliminate boundary artifacts and standardize the spatial dimensions across the dataset, the images were symmetrically cropped to a normalized field of view (170 px, corresponding to ~ 894.70 µm). Following the identification of the geometric centroid for each discrete fluid phase, the area moments of inertia were computed using the discrete pixel-wise summation

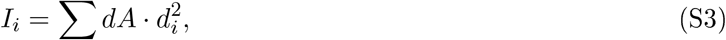

where *i* ∈ {*x, z}, dA* is the differential area element (corresponding to a single pixel), and *d*_*i*_ is the orthogonal distance from that pixel to the respective bending axis. The polar moment of inertia was subsequently derived using the perpendicular axis theorem,

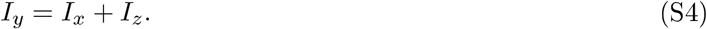

The heat maps presented in Figure 4Dii of the main text visually map these spatial distance distributions (*d*_*i*_) relative to the designated axes of rotation. Finally, the computed moments for each particle configuration were averaged across the sample population and converted from pixel coordinates to physical units (mm^4^).

## Note S9: Imaging the fibers’ cross-sectional profiles

To characterize the internal architecture of the extruded filaments, volumetric fluorescence data were acquired via confocal microscopy. The fabricated fibers were secured under mild tension within a custom 3D-printed alignment stage to prevent physical drift and ensure stable, high-resolution optical sectioning. Throughout the acquisition, the samples were continuously hydrated with deionized water to preserve the integrity of the hydrogel and optimize optical clarity. Longitudinal z-stacks were captured along the fiber axis, yielding localized 3D spatial representations of the embedded fluidic patterns. The resulting image stacks were subsequently processed using a custom Python routine that digitally resliced the volumetric data into a sequence of orthogonal 2D cross-sections. This workflow enabled the generation of continuous spatial video reconstructions, allowing fine-grained structural analysis of the sculpted internal topologies. To facilitate user interaction, a graphical user interface (GUI) was integrated into the core processing algorithm with the assistance of generative AI models (GitHub Copilot Student); the complete source code is publicly accessible via GitHub (https://github.com/masahinphd/actisculpt_confocal).

**Figure S8:**
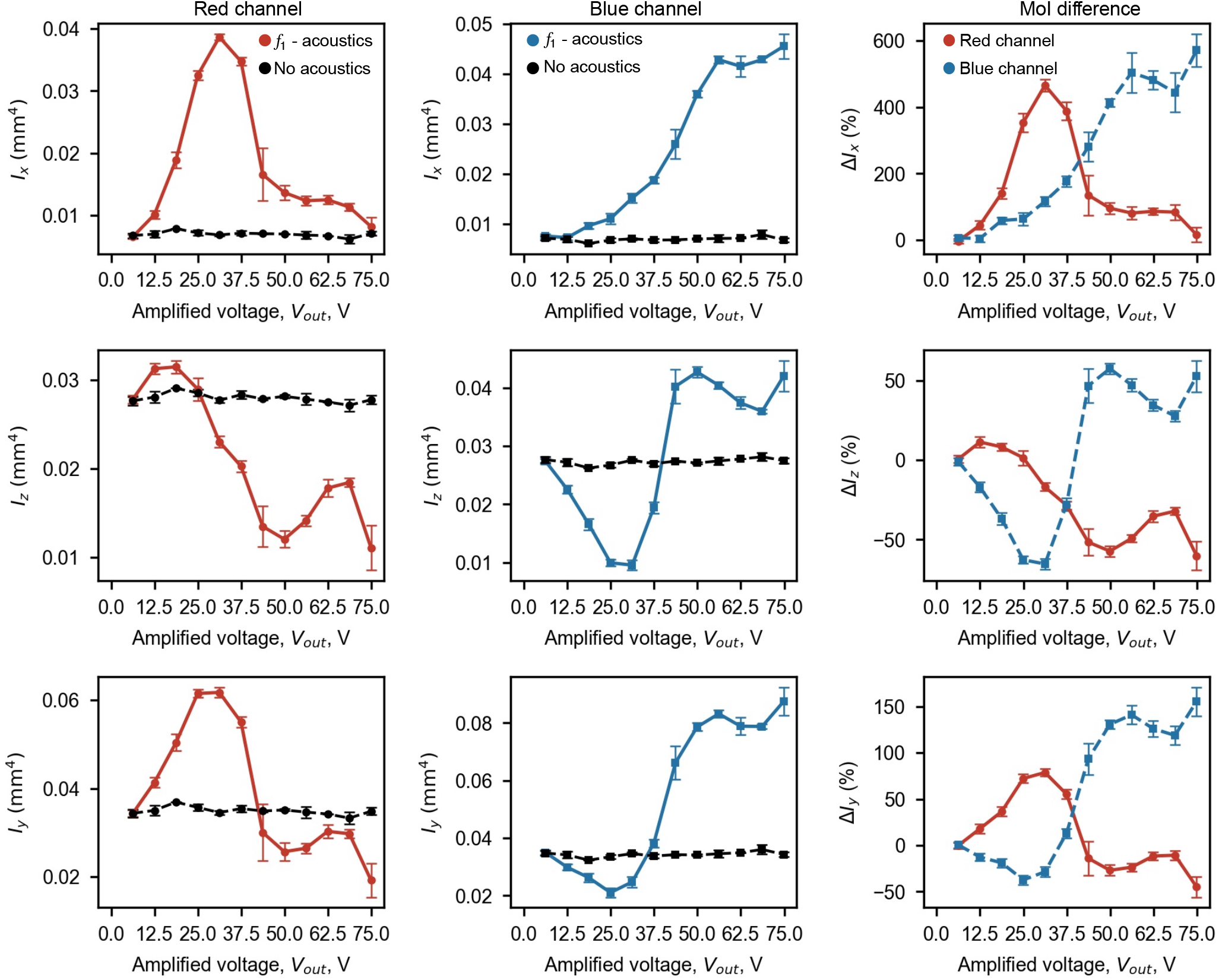
Modulation of area moments of inertia across principal axes under varying acoustic actuation of *f*_1_. Rows correspond to the x-, z-(polar), and y-axes of rotation. The first and second columns display the calculated moments for the blue and red fluorescent fluid domains, respectively, comparing the acoustically sculpted particles (red and blue traces) against the non-actuated reference baseline (black traces). The third column quantifies the relative percentage change in the moment of inertia for the actuated particles relative to the controls, illustrating the magnitude of mechanically tunable anisotropy achieved across the applied voltage sweep.

**Figure S9:**
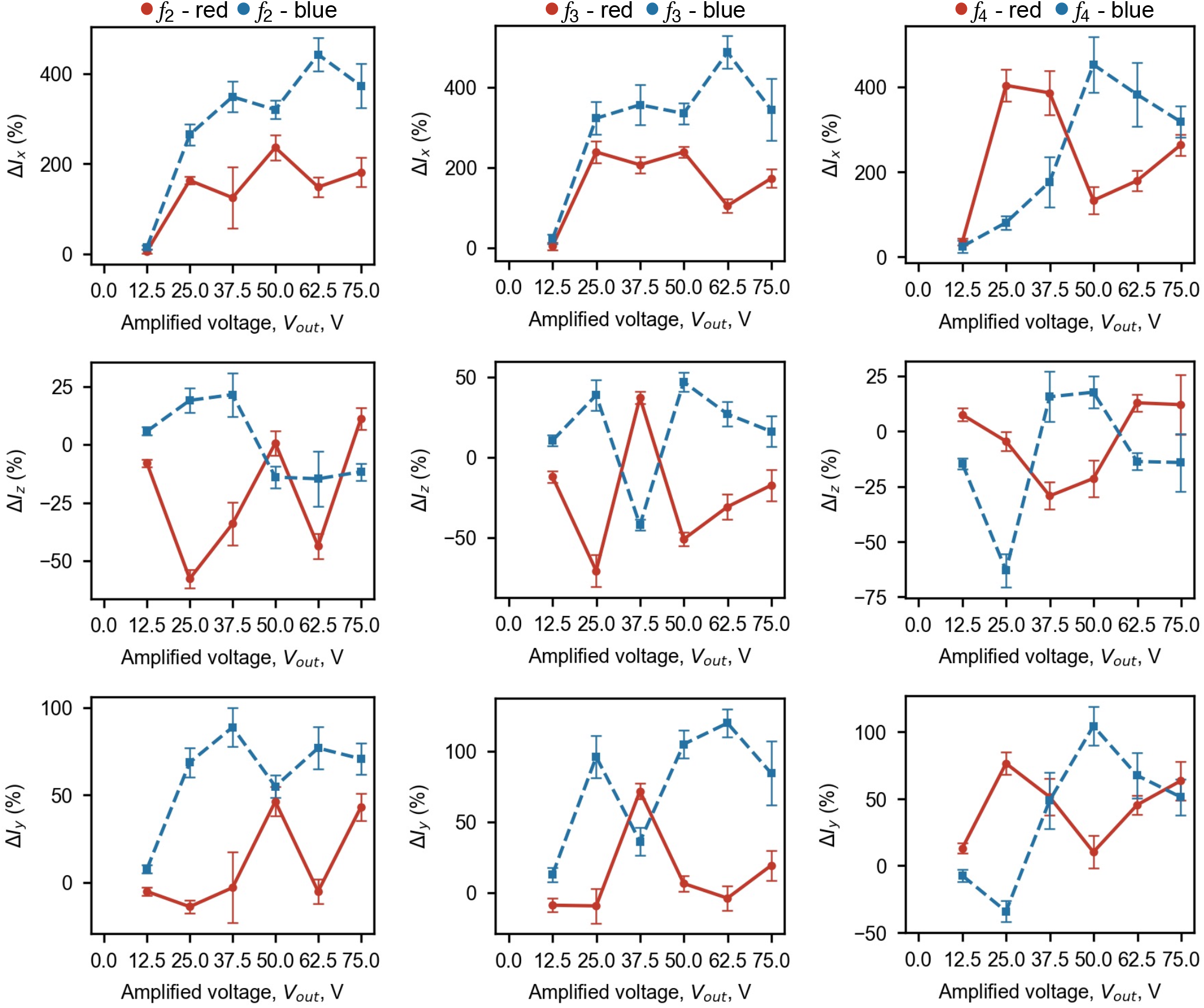
Extended relative percentage change of the moment of inertia over actuation sites *f*_2_, *f*_3_, and *f*_4_. Rows correspond to the x-, z-(polar), and y-axes of rotation. Columns correspond to the actuation sites *f*_2_, *f*_3_, and *f*_4_, respectively.

### Longitudinal spatial measurements and acoustic attenuation analysis

To verify the experimental cycle lengths against theoretical time-of-flight predictions, we analyzed the longitudinal intensity profiles extracted from the confocal scans (as depicted in Figure 5Biii of the main text and Movie S1). Corresponding analytical expected fiber lengths for one period are calculated by considering the real measured dimensions of the channel cross-section, 1038.21 *±* 29.23 µm and 1017.47 *±* 34.98 µm for the channel width and height, respectively, and a 250 ms total system signaling lag. The estimated lengths can then be calculated as follows:

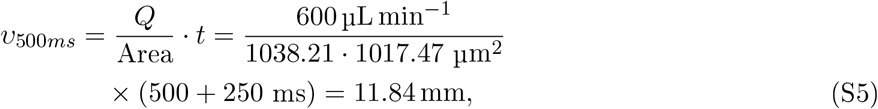

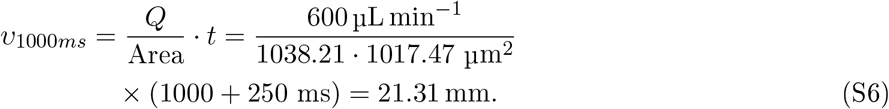

An important caveat for a velocity calculation that depends heavily on the channel cross-section is that the channel may expand during fiber extrusion, since it is partially closed by polymerized polymer. This observation is also supported by macroscopic measurements of fiber extrusion (see Note S10, below).

The spatial periodicity was measured by identifying the precise coordinates at which the fluorescence intensity begins to rise for consecutive *f*_2_ actuation peaks. At a constant *Q* = 600 µL min^−1^, the onset points for the 500 ms actuation cycle yielded a spatial period of 11.47 *±* 0.25 mm (*d <* 4%). Similarly, for the 1000 ms cycle, the respective onset points corresponded to an empirical cycle length of 19.20 *±* 0.57 mm (*d <* 10%) (Table S1).

**Table S1:** Qualification of the spatial periodicity via the initial-rise-point measurements of fibers created at 500 and 1000 ms actuation pulses of *f*_2_, and the combinatorial actuation of *f*_1_ and *f*_4_.

| Measurement | 500 ms<br>(mm) | 1000 ms<br>(mm) | Combinatorial<br>(mm) |
| --- | --- | --- | --- |
| Minimum $L$ | 11.21 | 18.54 | 7.93 |
| Maximum $L$ | 11.71 | 19.64 | 8.42 |
| Median $L$ | 11.47 | 19.40 | 8.17 |
| Average | 11.47 | 19.20 | 8.17 |
| Std. dev. | 0.25 | 0.57 | 0.24 |

For combinatorial actuation sequences, the spatial offset between distinct encoded features was quantified by tracking the initial rise of each feature’s intensity peak. To validate these empirical measurements, we compared the extracted spatial delay to the known geometric distance between the first (*f*_1_) and fourth (*f*_4_) interdigital transducers. The physical edge-to-edge separation between these IDTs is 10.02 mm. However, because the acoustic waves from both transducers propagate toward each other and attenuate through the fluid, the effective actuation distance is reduced by their respective attenuation lengths. The acoustic attenuation length (*α*^−1^) for each IDT was approximated using the relation

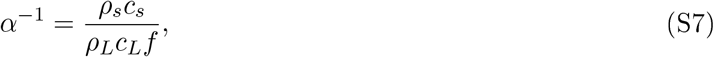

where *ρ*_*L*_ (1060 kg*/*m^3^) and *ρ*_*s*_ (4628 kg*/*m^3^) represent the densities, and *c*_*L*_ (1500 m*/*s) and *c*_*s*_ (3931 m*/*s) denote the speeds of sound for the liquid mixture and the LiNbO_3_ substrate, respectively. This analytical model assumes a simplified acoustic pathway, neglecting the negligible influence of the ultrathin PDMS membrane. Based on their specific resonant frequencies, the attenuation lengths were calculated as 1.22 mm for *f*_1_ and 0.96 mm for *f*_4_. Subtracting these attenuation zones from the physical transducer separation yields a theoretical effective distance of 7.85 mm (10.02 mm − 1.22 mm − 0.96 mm). This theoretical prediction closely aligns with the manually decoded spatial offset of 8.174 *±* 0.244 mm (*d <* 5%), derived from measured peak positions in Figure 5Ciii of the main text, where the selected points are represented in Table S1.

## Note S10: Macroscopic optical tracking of dynamic fiber extrusion

To characterize the spatial periodicity of fibers fabricated with extended actuation cycles, macroscopic optical recordings were continuously acquired during extrusion using a fixed-position CMOS camera (1080 *×* 1920 px, 30 fps). The precursor fluid comprised a vertically stacked bilayer, with the lower stream doped with a commercial blue colorant to provide high-contrast visual tracking of the acoustic deformation. Post-acquisition video processing was performed in ImageJ; sequences were trimmed to isolate steady-state extrusion regions containing a minimum of two complete acoustic modulation cycles. Spatial calibration was established using the known microchannel width, allowing both the axial cycle length (*L*_*cycle,theory*_) and the average experimental fiber extrusion velocity (*υ*_*exp,avg*_) to be quantified across the high-throughput flow regimes by measuring the longitudinal distance between consecutive sculpted phase boundaries. These measured values were subsequently compared against analytical predictions for a single pulse cycle length, yielding both idealized (*L*_*cycle,theory*_) and corrected (*L*_*cycle,corrected*_) theoretical estimates. The corrected analytical model specifically incorporated the empirical extrusion velocities and accounted for the experimentally determined 0.25 s total system lag. This comparative analysis confirmed that the elastomeric PDMS channel expands under the elevated backpressure induced by the polymerizing fiber. This volumetric expansion increases the effective cross-sectional area of the channel, causing the true extrusion velocity to deviate from idealized, pump-driven *Q* calculations (Table S2). As anticipated, this velocity divergence becomes more pronounced at elevated *Q*, where increased hydraulic resistance exacerbates the channel expansion.

**Table S2:**
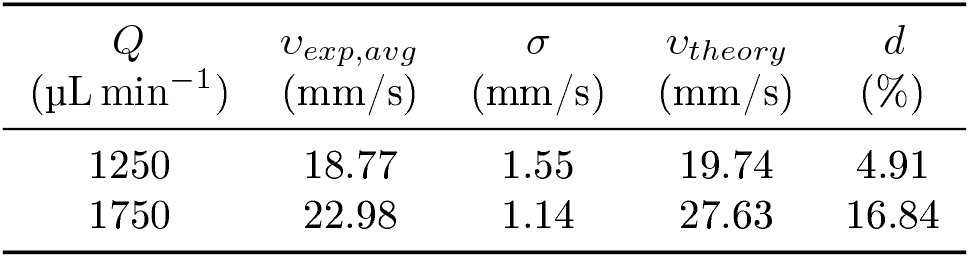
Experimental validation of the theoretical velocities, measuring the extrusion speed of the fiber out of the channel nozzle tip.

## Note S11: Tunable axial periodicity via pulse shaping

To assess the dynamic actuation limits of the continuous extrusion process in terms of final pattern convergence, we measured the spatial footprint of the transient fluidic responses, i.e., the ramping and linear deformation phases, under variable total actuation times (Figure S11A). During extrusion, a temporal lag dictates a transition phase until the final cross-sectional morphology is reached after the acoustic field is toggled. Because abrupt power switching induces transient hydrodynamic instabilities, we implemented a smoothed pulse-shaping protocol, *τ*_*actuation*_, featuring a gradual amplitude ramp, *τ*_*ramp*_ (Figure S10A). Combined with hardware communication latency, this introduces a total system delay, *τ*_*delay,total*_ = *τ*_*ramp*_ + *τ*_*latency*_, of approximately 0.25 s to 0.35 s. At *Q* = 600 µL min^−1^, confocal reconstructions of the extruded fibers allowed us to measure the physical lengths of these actuation phases for total pulse durations (*τ*_*actuation,total*_) of 0.5, 1, and 2 s. The measured actuation lengths, comprising the sum of the ramp-up and linear phases, scaled to 3.84, 6.71, and 11.90 mm, respectively (Figure S11B). While this relationship remains proportional for the 1 s and 2 s pulses, a slight deviation occurs at 0.5 s, where the system delay becomes comparable to the pulse duration. Furthermore, the spatial distance required for the fluid to “ramp” to its fully sculpted shape remained stable for the 1 s and 2 s pulses but contracted at 0.5 s. This contraction indicates that the fluid does not have sufficient time to converge to its target geometry, thereby defining the lower operational limit for stable feature encoding under these specific acoustic conditions.

Because confocal microscopy is inherently constrained by a limited field of view, we transitioned to macroscopic video tracking of food-dye-labeled streams to capture multiple consecutive deformation cycles. To define the limits of longitudinal patterning at higher *Q*, we investigated the system’s dynamic response under varying actuation duty cycles, shifting our focus from cross-sectional shape convergence to macroscopic spatial periodicity. We systematically varied the actuation duration (*τ*_*actuation*_) between 100 and 2000 ms. This corresponded to total cycle periods of 0.5 to 4.3 s (modulation frequencies of 0.23 to 2 Hz) during the extrusion of a partially dyed fiber under central actuation (*f*_2_, 320 mV) (Figure S10B, Movie S2). Macroscopic optical tracking of the extrudate revealed that the axial cycle length (*L*_*cycle*_) scales proportionally with the pulse duration but is not aligned with *Q* as determined by *L*_*cycle,theory*_ (Figure S11C). Correcting the time-of-flight calculation for the system lag and the measured extrusion velocity (*υ*_*exp,avg*_) yields the corrected length (*L*_*cycle,corrected*_), confirming that the PDMS channel expands while the fiber is extruding. Collectively, these results demonstrate that ActiSculpt provides robust temporal control over fiber morphology, enabling the on-demand scaling of cross-sectional features along the continuous filament.

**Figure S10:**
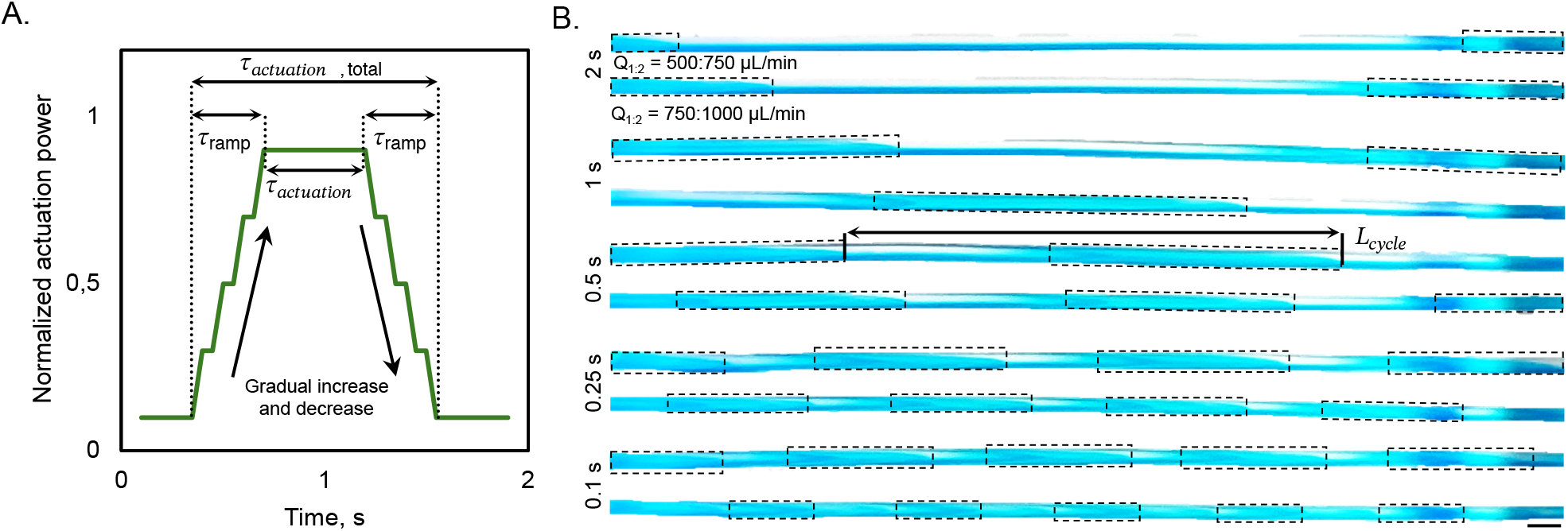
Temporal modulation of fiber cross-sections via programmable acoustic actuation. A. Schematic representation of the actuation profile with ramp-up (*τ*_*offset*_), actuation (*τ*_*actuation*_), and relaxation (*τ*_*relax*_) phases. A smooth sinusoidal signal profile was introduced to reduce abrupt transitions. B. Optical recordings of fiber extrusion under different *τ*_*actuation*_ settings and flow-rate conditions. The dyed fluid was used to visualize the dynamic sculpting of the cross-section.

**Figure S11:**
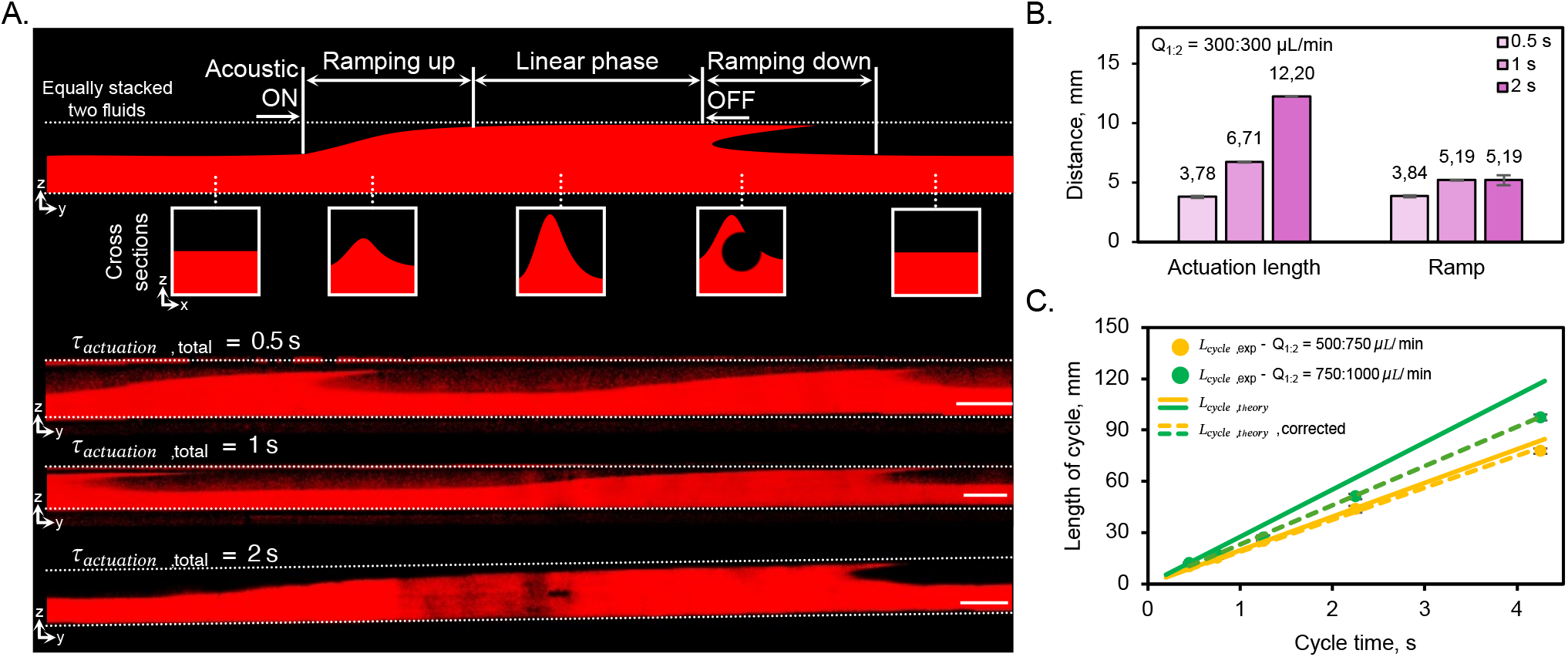
Temporal modulation of fiber cross-sections via dynamic acoustic actuation. A. Schematic representation of the actuation profile detailing the ramp-up, linear, and ramp-down phases, alongside confocal longitudinal reconstructions of fibers extruded under three distinct actuation durations. B. Quantification of the total actuation length (the sum of the ramp-up and linear phases) and the isolated ramp distance across varying pulse times; the data highlight the spatial footprint required for the fluid interface to converge to its target geometry. C. Macroscopic measurement of the axial cycle length (*L*_*cycle*_) extracted from optical recordings of continuous, food-dye-labeled streams. The plot compares experimental spatial periodicities against theoretical predictions across varying flow rates and duty cycles.

## Movie Captions

**Movie S1**. Confocal reconstruction of pulsed-actuation fibers showing dynamic encoding of longitudinal morphology under a single transducer (*f*_2_) actuated in 500 ms/1000 ms on-off cycles. The movie corresponds to Figure 5B.

**Movie S2**. Macroscopic optical recording of continuous fiber extrusion under varying actuation duty cycles, illustrating spatial periodicity scaling with pulse duration. The movie corresponds to Figure S10B.

**Movie S3**. Recording of the ActiSculpt nozzle printing process on the motion-controlled 3D printer, including zoomed-out and zoomed-in views of the extruded fiber. The movie corresponds to Figure 6B.

